# Dynamics of the *Bacillus subtilis* Min system

**DOI:** 10.1101/2020.04.29.068676

**Authors:** Helge Feddersen, Laeschkir Würthner, Erwin Frey, Marc Bramkamp

**Affiliations:** Institute for General Microbiology, Christian-Albrechts-University Kiel, Am Botanischen Garten 1-9, 24118 Kiel, Germany; Arnold-Sommerfeld-Center for Theoretical Physics and Center for NanoScience, Ludwig-Maximilians-Universität München, Theresienstr. 37, 80333 Munich, Germany; Ludwig-Maximilians-Universität München, Faculty of Biology, Großhaderner Straße 2-4, 82152 Planegg-Martinsried, Germany

**Keywords:** *B. subtilis*, Min system, cell division, FRAP, PALM, super resolution microscopy, protein patterns, reaction diffusion equations

## Abstract

Division site selection is a vital process to ensure generation of viable offspring. In many rod-shaped bacteria a dynamic protein system, termed the Min system, acts as a central regulator of division site placement. The Min system is best studied in *Escherichia coli* where it shows a remarkable oscillation from pole to pole with a time-averaged density minimum at midcell. Several components of the Min system are conserved in the Gram-positive model organism *Bacillus subtilis*. However, in *B. subtilis* it is believed that the system forms a stationary bipolar gradient from the cell poles to midcell. Here, we show that the Min system of *B. subtilis* localizes dynamically to active sites of division, often organized in clusters. We provide physical modelling using measured diffusion constants that describe the observed enrichment of the Min system at the septum. Modelling suggests that the observed localization pattern of Min proteins corresponds to a dynamic equilibrium state. Our data provide evidence for the importance of ongoing septation for the Min dynamics, consistent with a major role of the Min system to control active division sites, but not cell pole areas.

## Introduction

The spatiotemporal regulation of cell division in bacteria is an essential mechanism ensuring correct partitioning of DNA to produce viable daughter cells upon division. The best studied model organisms in this aspect are the rod-shaped Gram-positive and Gram-negative bacteria *Bacillus subtilis* and *Escherichia coli*, respectively. Both species divide precisely at the geometric middle via binary fission. The earliest observed event in this process is the formation of the Z-ring, a ring-like structure consisting of polymerized FtsZ proteins, a highly conserved homologue of eukaryotic tubulin [1-5]. Once assembled, FtsZ acts as a dynamic scaffold and recruits a diverse set of proteins forming the divisome, a complex that mediates cytokinesis [6-8]. Recently, treadmilling of FtsZ filaments was shown to drive circumferential peptidoglycan (PG) synthesis [9]. However, it is still not fully understood how FtsZ finds the precise midplane of the cell. In *E. coli* and *B. subtilis* the nucleoid occlusion (NO) and the Min system, two negative FtsZ regulators, have been shown to confine its action spatially to the center of the cell. Noc in *B. subtilis* and SlmA in *E. coli* bind to DNA and inhibit FtsZ polymerization across the nucleoid [10-14]

The Min system in *E. coli* consists of the three proteins MinC, MinD and MinE [15, 16], and has been studied extensively both experimentally [17-29] and theoretically [29-35]. MinC is the inhibitor of Z-ring formation, inhibiting the bundling of FtsZ filaments [22, 36-39]. MinC is localized through MinD, a protein that belongs to the WACA (Walker A cytomotive ATPase) family [40, 41]. Upon binding ATP, MinD dimerizes and associates with the membrane through a conserved C-terminal membrane targeting sequence (MTS) [3, 42, 43]. MinC and MinD were described to form large ATP-dependent alternating polymers that assemble cooperatively and locally inhibit FtsZ bundling [44, 45]. In absence of MinCD, cells frequently produce the name giving anucleate minicells [46, 47]. The *E. coli* MinCD complexes are disassembled and detached from the membrane by MinE, a protein that forms a ring-like density profile at the rim of MinD assemblies [48, 49] and binds to the membrane via an amphipathic helix serving as membrane targeting sequence (MTS) [50, 51]. MinE triggers ATPase activity of MinD, leading to membrane detachment of MinCD [27]. Cytosolic MinD rebinds ATP and binds the membrane again, thereby leading to a remarkably robust oscillation of the Min system in *E. coli* [25, 27, 52, 53]. Min protein dynamics are a paradigmatic example of cellular self-organization [54]. Due to the simplicity of the system, it has been subject to several molecular modelling studies and in vitro reconstructions [26-35].

The Min system in *B. subtilis* lacks MinE as essential factor that is responsible for Min oscillation in *E. coli*. In contrast, Min proteins do not oscillate in *B. subtilis* but are believed to form a stationary bipolar gradient decreasing towards midcell, therefore restricting assembly of a functional FtsZ ring to the midplane of the cell [3, 8]. The spatial cue for a gradient in *B. subtilis* is provided by DivIVA [55, 56]. DivIVA targets and localizes to negatively curved membrane regions [57]. MinJ acts as a molecular bridge between MinD and DivIVA [58, 59]. MinJ contains six predicted transmembrane helices and a PDZ domain, which often participate in protein-protein interactions [60]. Due to the polar targeting of DivIVA, MinCDJ should form a stationary polar gradient decreasing towards midcell, restricting FtsZ polymerization spatially at midcell. However, several studies suggest that the *B. subtilis* Min system may rather act downstream of FtsZ ring formation by preventing re-initiation of division at former sites of cytokinesis [58, 59, 61].

We have recently analyzed DivIVA dynamics in *B. subtilis* and found that Min proteins redistribute from the cell poles to midcell as soon as a septum is formed [62], which prompted us to reanalyze Min protein dynamics in this organism. To this end, we generated a set of new fusions to DivIVA, MinD and MinJ. To avoid overexpression artifacts that would corrupt protein dynamics studies, we generated strains where the native gene copies were replaced by functional fluorescent fusions. These allelic replacements were used to determine precise molecule counts per cell. Using fluorescent recovery after photo-bleaching (FRAP) we determined protein dynamics of the individual Min proteins. We then calculated protein diffusion coefficients that were further used for modeling and simulations of the observed Min dynamics. We finally analyzed the nanoscale spatial distribution of the Min proteins in *B. subtilis* by single molecule localization microscopy (SMLM). Our data are consistent with a dynamic turnover of MinD between membrane and cytosol. Moreover, our SMLM data support a model in which the Min complex is in a dynamic steady state that is able to relocalize from the cell pole to the septum facilitated by a geometric cue, namely the invagination of the membrane at the septum. Based on our experimental data, we propose a minimal theoretical model for the Min dynamics in *B. subtilis* in realistic 3D cell geometry. The model is based on a reaction-diffusion system for MinD and incorporates the effects of DivIVA and MinJ implicitly through space-dependent recruitment and detachment processes. Our computational analysis of the mathematical model reproduces qualitative features of the Min dynamics in *B. subtilis* and shows that localization of MinD to the poles or septum corresponds to a dynamic equilibrium state. Furthermore, our model suggests that a geometric effect alone could explain septum localization of MinD once DivIVA is recruited to the growing septum, therefore highlighting the importance of geometry effects that cannot be captured in a simplified one-dimensional (1D) model.

## Results

### Construction of fluorescent fusions with native expression level

Even though the Min system in *B. subtilis* has been extensively investigated before, most studies were conducted using strains that overexpress fluorescent fusions from ectopic locations upon artificial induction [63, 64], leading to non-native expression levels that can alter the native behavior of fine-tuned systems like the Min system. Additionally, even small populations of a protein from overexpression make it difficult to identify a dynamic fraction through diffraction limited microscopy [65]. Hence, we aimed to re-characterize the dynamics of the Min components in *B. subtilis* using strains that avoid or minimize overexpression artifacts and, hence, created a set of allelic replacements (**Fig. S1, Fig. S2**).

Dysfunctionality or deletion of Min components in *B. subtilis* manifests in an easily observable phenotype of increased cell length and DNA free minicells (**Tab. 1**). This allows rapid evaluation of the functionality of fluorescent-fusions in the constructed strains by comparing cell length and number of minicells between mutant and the wild-type strain (**Tab. 1**).

**Tab. 1:** Phenotypic characterization of relevant strains. For determination of the growth rate constant µ the optical density at 600nm of exponentially growing cells was measured. Cell lengths and the ratio of minicells were determined microscopically using Fiji, with n ≥ 200. Strains were grown in independent triplicates, with differences reflected in the standard deviation.

| Strain | Description of strain | Mean growth rate | Mean cell length | Minicells |
| --- | --- | --- | --- | --- |
| | | constant [$\mu$] | [$\mu\text{m}$] | [%] |
| 168 | Wild type | $0.53 \pm 0.004$ | $3.106 \pm 0.773$ | 0.3 |
| 3309 | $\Delta\text{minCD}$ | $0.45 \pm 0.021$ | $7.639 \pm 2.702$ | 45.8 |
| RD021 | $\Delta\text{minJ}$ | $0.51 \pm 0.049$ | $6.645 \pm 2.022$ | 13.8 |
| 4041 | $\Delta\text{divIVA}$ | $0.46 \pm 0.020$ | $8.128 \pm 3.404$ | 29.6 |
| BHF011 | Dendra2-MinD | $0.49 \pm 0.004$ | $2.664 \pm 0.607$ | 0.9 |
| BHF017 | msfGFP-MinD | $0.55 \pm 0.004$ | $4.221 \pm 1.036$ | 9.1 |
| JB38 | MinJ-Dendra2 | $0.51 \pm 0.006$ | $3.437 \pm 1.056$ | 0 |
| BHF007 | MinJ-msfGFP | $0.57 \pm 0.013$ | $3.375 \pm 0.755$ | 0.3 |
| JB40 | MinJ-mNeonGreen | $0.57 \pm 0.002$ | $3.156 \pm 0.673$ | 0 |
| JB36 | DivIVA-Dendra2 | $0.50 \pm 0.007$ | $4.327 \pm 0.917$ | 8.0 |
| **1803** | DivIVA-GFP | $0.45 \pm 0.021$ | $3.309 \pm 0.733$ | 1.1 |
| **BHF028** | DivIVA-mNeonGreen | $0.54 \pm 0.029$ | $5.419 \pm 1.351$ | 5.3 |
| **JB37** | DivIVA-PAmCherry | $0.51 \pm 0.019$ | $4.346 \pm 1.105$ | 3.3 |

Here, we generated functional fusions to MinD (Dendra2 [66]) and MinJ (msfGFP [67] and mNeonGreen [68]) as judged by cell length, number of minicells and subcellular protein localization (**Fig. S2, Tab. 1**). Dendra2-MinD displayed a phenotype comparable to the wild-type, but could unfortunately not be used for FRAP studies due to the fluorophore characteristics. Therefore, another strain expressing msfGFP fused to MinD was created. This fusion protein was largely functional according to cell length and number of minicells (**Fig. S2, Tab. 1**). When **msfGFP-minD** was transformed in a genetic background of *ΔminJ* or *ΔdivIVA*, the fluorescent signal was, as expected, distributed in the cytosol, sometimes forming small foci. MinJ-msfGFP also lost its polar and septal localization upon deletion of *divIVA*, as it has been reported before [58]. Hence, these strains were not used for further analysis of protein dynamics.

When constructing DivIVA fluorescent fusions, several different fluorophores (FPs) were successfully fused to DivIVA, namely mCherry2, mNeonGreen, Dendra2, PAmCherry, mGeosM and Dronpa [66, 68-72], with linkers between 2 and 15 amino acids. Unfortunately, all of them showed a mild or strong phenotype, some even severe protein mislocalization, hinting towards limited functionality of these DivIVA fusion proteins ([73] and data not shown). Since this did not meet the set standards for this study, we turned towards strain 1803 [74], carrying a *divIVA-GFP* copy with its native promoter in the ectopic *amyE* locus. While *DivIVA-GFP* has been shown to not fully complement a *ΔdivIVA* strain [74, 75], it still localizes correctly and can be used for studies of DivIVA dynamics [62, 75]. Additionally, we performed FRAP on DivIVA-mNeonGreen, which only shows a mild phenotype (**Tab. 1**), in wild type and Min knockout backgrounds to be able to compare it with the effect of the extra copy of DivIVA in strain 1803 (**Fig. S3**).

All fluorescent fusions were analyzed via SDS-PAGE with subsequent visualization through in-gel fluorescence or western blotting (**Fig. S4**. We used in-gel fluorescence to obtain estimations about the number of molecules of the Min proteins during mid-exponential phase. We calculated protein numbers relative to the total amount of MinD that was quantified under same growth conditions using mass spectrometry before [76] (**Tab. S1, Fig. S5**). MinD proteins are highly abundant (3544 proteins per cell), while DivIVA numbers are less than 50% of that (1690 proteins per cell). MinJ has only 16% of MinD abundancy (576 proteins per cell).

### The Min system in *B. subtilis* is in a dynamic steady state

Strains expressing functional Min fusions were then used for microscopic analysis of protein dynamics using fluorescent recovery after photobleaching (FRAP) experiments. Surprisingly, all three components of the Min system showed relatively fast diffusion in FRAP (**Error! Reference source not found., Figure 2, Tab. 2**). Unfortunately, Dendra2-MinD could not be used for FRAP studies, because excitation with 488 nm leads to a significant green to red conversion during the course of the experiment. When converting all proteins from green to red prior the FRAP experiment with UV light (405 nm), the red fluorescent signal was poor and bleaching of most proteins occurred during the first image acquisitions, prohibiting reliable quantification. Upon converting protein locally at one of the poles or a septum with a short laser pulse at 405 nm and subsequent imaging in the red channel, very fast diffusion of converted Dendra2-MinD throughout the cell could be observed (data not shown). However, the signal was too dim to be quantified satisfactorily. Therefore, we utilized the strain expressing msfGFP-MinD (BHF017) and observed a fast fluorescence recovery (T_1/2_= 7.55s), indicating rapid exchange of MinD molecules around the division septum, similar to what was previously reported for MinC [65]. Bleaching of MinD at a septum was very efficient (**Error! Reference source not found. a, upper panel**), the exchange of MinD molecules at the bleached spot appeared to include proteins localized distant from the bleached septum as well as in the vicinity, since the fluorescent signal in the cell reduced evenly over the whole cell length during recovery. Furthermore, around 79% of the msfGFP-MinD population appeared to be mobile (**Figure 2, Tab. 2**).

**Tab. 2:** Results of FRAP analysis for the Min proteins in different genetic backgrounds.

| Protein and genetic background | Fluorophore | Diffusion coefficient<br>[$\mu\text{m}^2 * 10^{-3} * \text{s}^{-1}$] | Half-Time<br>Recovery [s] | Mobile<br>Fraction |
| --- | --- | --- | --- | --- |
| **MinD in wild type** | msfGFP | 57.78 ± 10.05 | 7.55 ± 1.31 | 0.79 ± 0.06 |
| **MinJ in wild type** | msfGFP | 7.19 ± 2.27 | 62.35 ± 19.71 | 0.77 ± 0.14 |
| **MinJ in $\Delta\text{minCD}$** | msfGFP | 14.46 ± 9.54 | 30.18 ± 19.92 | 0.75 ± 0.07 |
| **DivIVA in wild type** | GFP | 3.39 ± 0.82 | 128.64 ± 30.92 | 0.65 ± 0.23 |
| **DivIVA in $\Delta\text{minCD}$** | GFP | 3.74 ± 1.36 | 116.79 ± 42.40 | 0.68 ± 0.26 |
| **DivIVA in $\Delta\text{minJ}$** | GFP | 8.57 ± 4.43 | 50.94 ± 26.35 | 0.49 ± 0.15 |
| **DivIVA in $\Delta\text{minCDJ}$** | GFP | 4.98 ± 2.93 | 87.67 ± 51.55 | 0.61 ± 0.27 |
| **DivIVA in wild type** | mNeonGreen | 7.23 ± 1.99 | 60.34 ± 16.57 | 0.64 ± 0.23 |
| **DivIVA in $\Delta\text{minCD}$** | mNeonGreen | 6.88 ± 2.76 | 63.43 ± 25.41 | 0.67 ± 0.20 |
| **DivIVA in $\Delta\text{minJ}$** | mNeonGreen | 18.01 ± 3.22 | 24.40 ± 4.33 | 0.39 ± 0.11 |
| **DivIVA in $\Delta\text{minCDJ}$** | mNeonGreen | 9.47 ± 4.26 | 46.07 ± 20.71 | 0.66 ± 0.23 |

**Figure 1:**
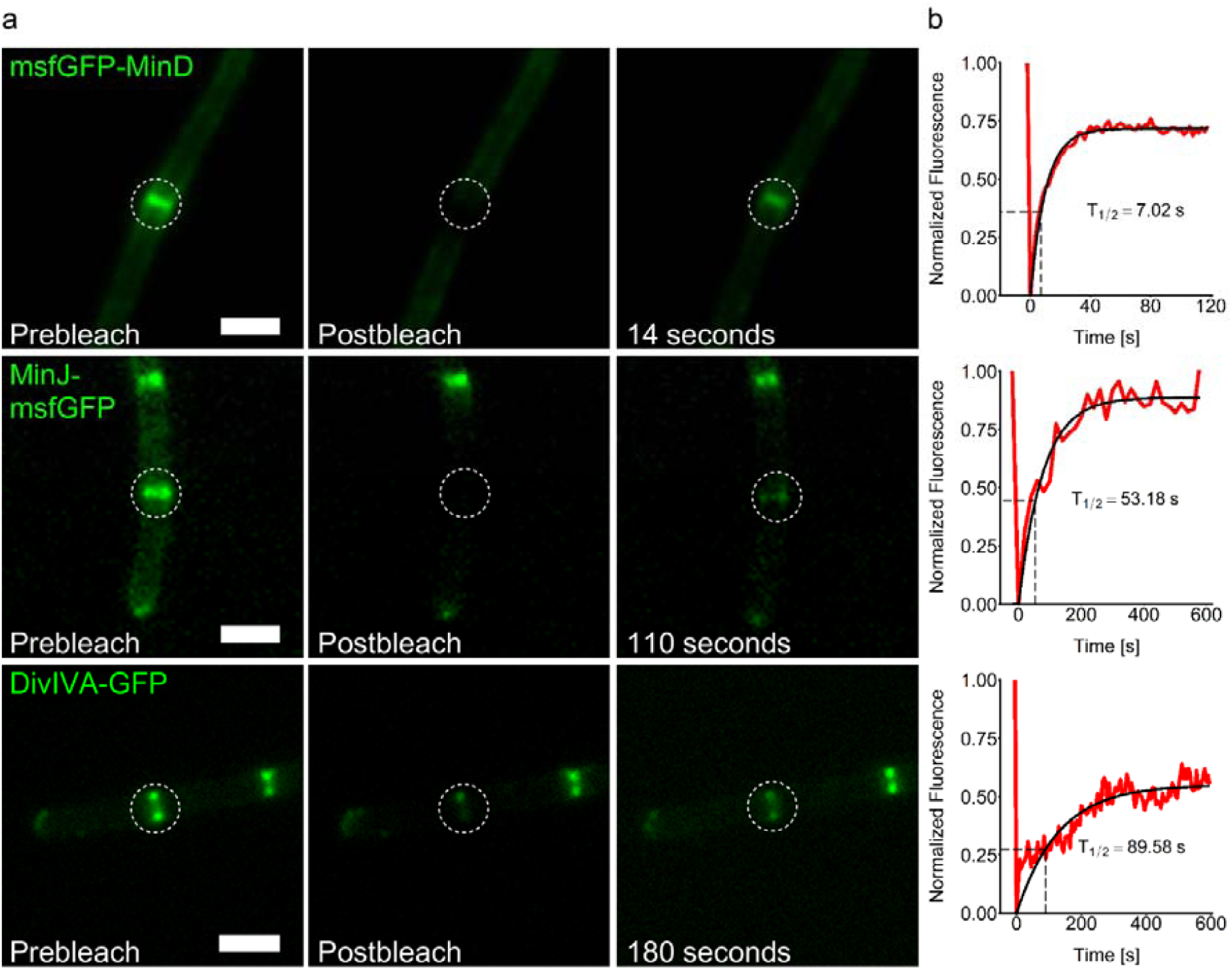
FRAP experiments in growing *B. subtilis* cells reveal Min protein dynamics. **(a)** Representative microscopy images of msfGFP-MinD (BHF017), MinJ-msfGFP (BHF007) and DivIVA-GFP (1803) before bleaching the indicated spot with a 488 nm laser pulse, directly after bleaching and after recovery of fluorescence. Scale bars 2 µm. (**b**) Representation of the normalized fluorescence recovery in the green channel over time. T_1/2_ = time when fluorescence recovery reaches half height of total recovery, indicated on the graph with a dashed square. The red line represents measured values, black the fitted values. Values were obtained as described in the methods **(Eq. 1-3)**.

**Figure 2:**
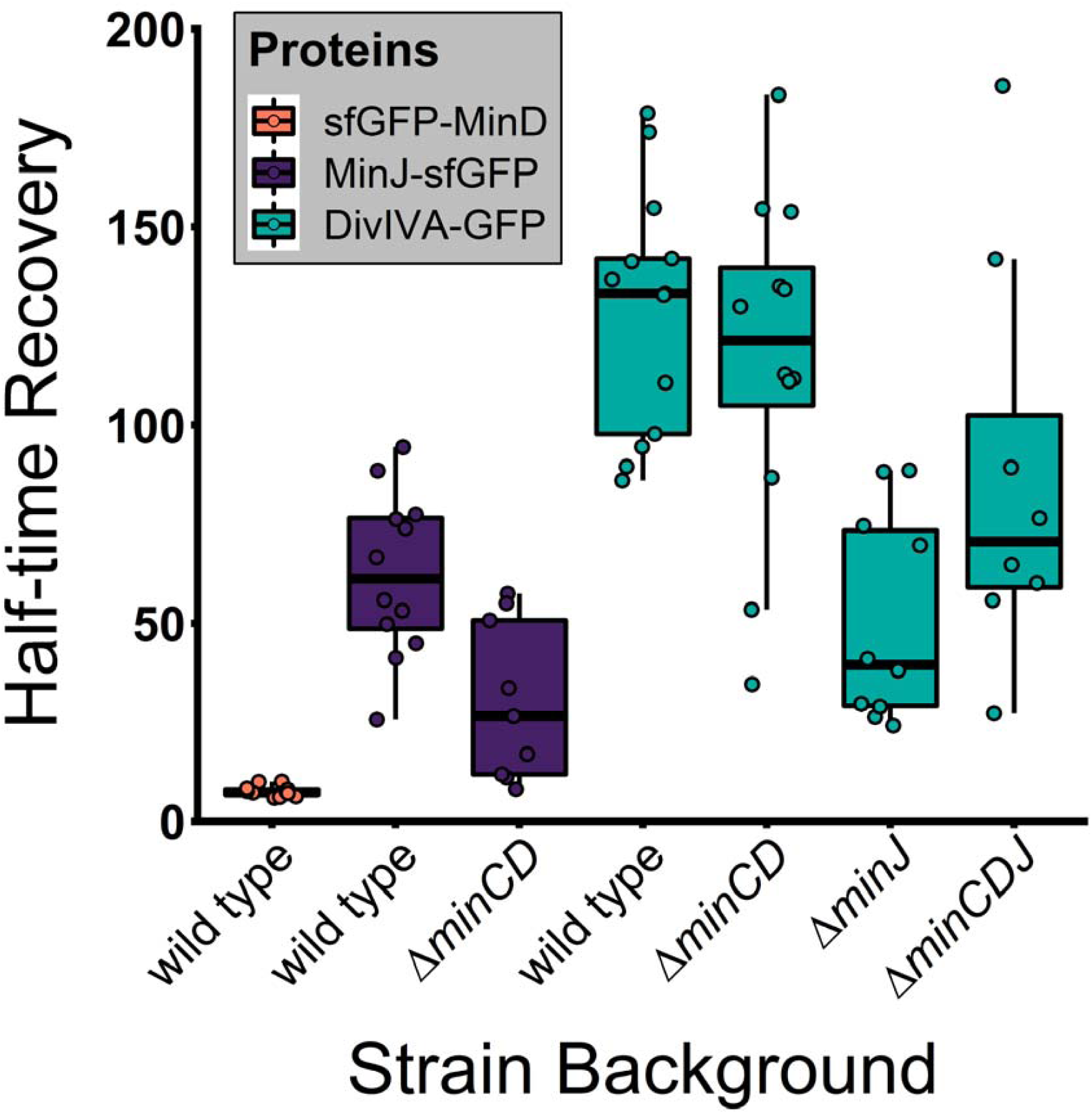
*B. subtilis* Min proteins form dynamic complexes. Shown are half-time recovery median indicated by black bar inside each box. Each box represents a different strain, also see **Tab. 2** for mean values. Every dot represents a single FRAP experiment, n ≥ 8.

Next, we investigated MinJ-msfGFP fluorescence recovery, which was considerably slower compared to msfGFP-MinD, but still indicating protein diffusion (T_1/2_ = 62.35s). MinJ contains six predicted transmembrane helices and therefore, a slower recovery was expected. Again, most of the MinJ-msfGFP protein pool appeared to participate in the fluorescence recovery (77%). When we measured *DivIVA-GFP* and DivIVA-mNeonGreen dynamics at septal localizations using FRAP, we observed similar mobility (DivIVA GFP T_1/2_ = 128.64s and DivIVA-mNeonGreen T_1/2_ = 60.34s). Since the DivIVA-GFP expressing strain has an extra copy of *divIVA*, it seems logical that the recovery time roughly doubles compared to the DivIVA-mNeonGreen expressing strain with only one copy of the gene. DivIVA has previously been reported as static [75], however those FRAP experiments were carried out using overexpression strains and a much shorter timeframe than here. Earlier observation from our own lab using a merodiploid strain already suggested that DivIVA is dynamic [62]. Roughly, two thirds of DivIVA molecules were participating in dynamics.

### Interaction of Min proteins influences their dynamics

To obtain a better understanding of the interactions between Min proteins and to find an explanation for the observed dynamics, we performed FRAP experiments in various genetic knockout backgrounds of Min genes. The Min system is hierarchically assembled with DivIVA recruiting MinJ, which then recruits MinD [58]. In agreement with that, we saw a loss of polar and septal msfGFP-MinD localization (BHF025 & BHF026) when we knocked out *minJ* or *divIVA* (data not shown). The same was true for MinJ-msfGFP upon knocking out *divIVA* (BHF032), further corroborating that DivIVA/MinJ complexes are required for controlled MinD localization. Therefore, we did not include these strains in the FRAP analysis. When *minCD* was knocked out in a strain expressing MinJ-msfGFP, the half-time recovery in FRAP dropped from 62s to 30s (**Figure 2, Tab. 2, Fig. S6**). This behavior indicates direct interaction between the two proteins. We cannot exclude that the phenotype itself impacts the dynamic behavior of MinJ, since cells are elongated and often re-divide after successful cytokinesis [61]. When *minCD* was knocked out in a *DivIVA-GFP* expressing strain (BHF040) however, we could not see any significant difference in fluorescence recovery. Since there is no direct interaction, DivIVA dynamics do not seem to be affected by MinCD directly nor indirectly, which includes the effects of the phenotype of elongated cells. Contrary to that, knocking out minJ sped up recovery of *DivIVA-GFP* (BHF041) significantly, with a recovery time less than half of the wild type, which was also true for DivIVA-mNeonGreen (BHF027) (**Tab. 2, Fig. S6**). This is consistent with a direct DivIVA MinJ interaction. Interestingly, there was also an impact on the mobile fraction, which decreased from around two thirds to roughly 40%-50% in both strains. Thus, dynamics are modulated by complex formation reflecting the expected protein hierarchy. MinD recruitment to midcell is fully dependent on DivIVA/MinJ. Since these proteins are only relocating to late stages of septum development, e.g. after a crosswall has started to form, we argue that this geometric change in the cell is important to redistribute MinD from the poles to midcell and establish a new dynamic steady state at the septum/new pole. This localization of MinD at midcell is lost if either DivIVA or MinJ are deleted, or MinD ATPase activity is abolished, as it can be observed in the G12V and K16A ATPase mutants of MinD [77]. Thus, maintenance of a steady gradient requires ATPase activity and is therefore similar to the *E. coli* system. Therefore, we aimed to support this hypothesis by mathematical modelling to understand the observed dynamics further.

### Theoretical model for MinD dynamics in *B. subtilis*

Previous theoretical studies of the Min system in *B. subtilis* are sparse. To our knowledge, there is actually only a single theoretical study that has investigated a mechanism for the polar localization of proteins [78]. In this work, the coupled dynamics of DivIVA and MinD are modeled by a reaction-diffusion system in one spatial dimension. Both MinD and DivIVA are considered to diffuse on the membrane and in the cytosol, and cycle between these two compartments by attachment and detachment processes. Membrane-bound MinD is assumed to be stabilized through DivIVA, hence its role is quite different from MinE, which destabilizes membrane-bound MinD. Moreover, it was argued that DivIVA requires the presence of MinD for membrane binding [78]; specifically, that DivIVA binds to and then stabilizes the edges of MinD clusters. Since the model was studied in one spatial dimension, the author accounted for geometric effects only implicitly by reducing the MinD attachment rate near the cell poles. The importance of ATP binding and hydrolysis on MinD activity has been discussed, but disregarded in the model as explicit coupling between cytosol and membrane (bulk-boundary coupling) was not considered. In summary, the model was a first and important theoretical analysis dissecting the relative role of MinD and DivIVA as well as their interplay in shaping protein localization in *B. subtilis*.

Here, on the basis of previous theoretical studies of intracellular protein dynamics [30, 32, 34, 79], we propose a minimal reaction-diffusion system to model Min localization in *B. subtilis*. Building on the idea of geometry sensing put forward in Ref. [79], our model provides a possible mechanism for how proteins sense cell geometry. This mathematical analysis shows that Min polarization and localization is established through a highly dynamic process driven by the ATPase activity of MinD. This implies Min protein gradients are maintained by genuine nonequilbrium processes and not by thermodynamic binding (chemical equilibrium) of Min proteins to a DivIVA template at the cell poles [3, 8].

We study protein dynamics in realistic three-dimensional (3D) cellular geometry, where proteins cycle between cytosol and membrane, and MinD diffuses with diffusion constants, *D*_*D*_ = 16 *µm*^2^/*s* and, *D*_*d*_ = 0.06 *µm*^2^/*s*, in the cytosol and on the membrane, respectively. Experimental studies have shown that DivIVA binds preferentially to regions of high negative membrane curvature and that MinJ localization is dependent on the presence of DivIVA [57, 58]. Moreover, MinJ is known to act as an intermediary between DivIVA and MinD [58]. Integrating this information into a mechanistic theoretical model, we consider fully resolved dynamics of MinD (including its ATPase cycle). The biochemical reaction scheme, illustrated in **Figure 3 a** and **b**, is based on the following molecular processes: (i) Attachment to and detachment from the membrane with rates *k*_*D*_ = 0.068 *µm*/*s* and 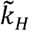 = 0.1 *s*^−1^ respectively. (ii) A nonlinear recruitment process of cytosolic MinD by membrane-bound MinD with rate 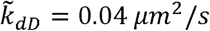 (iii) After detachment from the membrane, MinD is in an ADP-bound state and can rebind to the membrane only after nucleotide exchange which occurs at rate *λ* = 6 *s*^−1^. The protein numbers and membrane diffusion of MinD were extracted from our measurements (see **Tab. 2** and **Supplementary Material, Tab. S1 and Tab. S2**) and the values for the rate constants were estimated from previous work on protein pattern formation [30, 32, 34, 78]. The action on DivIVA and MinJ is accounted for effectively through space-dependent recruitment and detachment rates of MinD at membrane areas with a negative curvature; for details please refer to the Methods section (**Fig. S7**). Specifically, in our computational studies, using finite element methods (see Method section), it is assumed that in cells with no pre-existing division apparatus, MinD attachment at the poles is favored due to additional recruitment by MinJ (larger recruitment rate) and that detachment is reduced through stabilization by MinJ-DivIVA complexes (smaller detachment rate). The latter assumption is motivated by experiments, which suggest that DivIVA serves as a scaffold protein [57, 80]. Under the above conditions, MinD accumulates at both cell poles in a dynamic equilibrium with proteins constantly cycling between cytosol and membrane (**Figure 3 c**). In contrast, in the absence of preferential attachment at the cell poles facilitated by MinJ-DivIVA complexes, polar localization of MinD becomes unstable and the proteins become preferentially localized in the cell center (again in a dynamic equilibrium state). As can be inferred from **Figure 3 c**, this localization is, however, rather diffuse and covers the whole cell body. Next, we tested whether MinD can be localized at midplane in the presence of MinJ-DivIVA complexes once a septum has formed there. Indeed, emulating the presence of these complexes by an enhanced recruitment and detachment rate localized at the septum, our simulations show that MinD becomes sharply localized at midplane following the transfer of MinJ-DivIVA complexes from the poles to the septum (**Figure 3 b**). The width of the MinD distribution at midcell is determined by the interplay between membrane diffusion and localized recruitment of MinD at the septum (see Method section).

**Figure 3:**
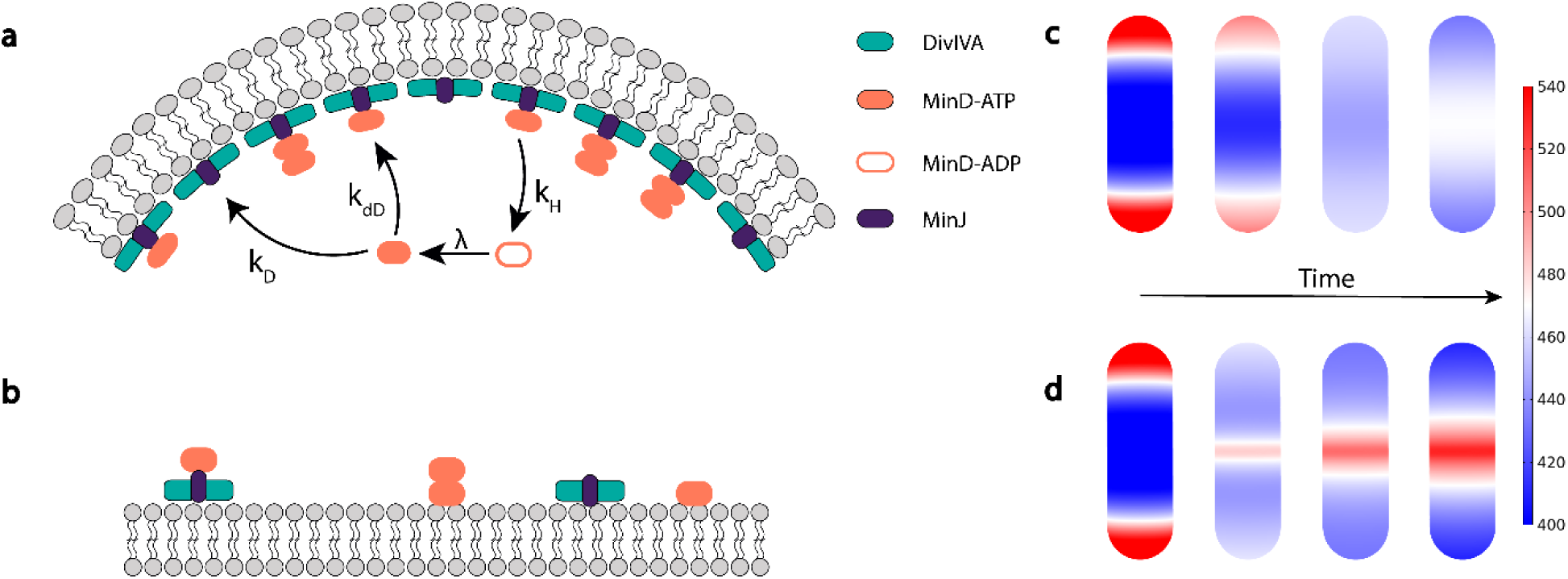
Model and simulation of the Min system in *B. subtilis*. (**a**) The geometry sensing protein DivIVA (green) preferentially localizes to regions of highest negative curvature and stabilizes MinJ (purple) to these regions. Membrane-bound DivIVA acts as a template for MinD-recruitment of cytosolic MinD-ATP (orange) facilitated through MinJ. MinD-ATP binds to the membrane with a rate k_D_ and recruits cytosolic MinD-ATP with a (space-dependent) recruitment rate k_dD_ to the membrane. Membrane-bound MinD is stabilized by MinJ-DivIVA complexes, which is reflected in a space-dependent detachment rate k_det_. After detachment, MinD is in a hydrolysed state MinD-ADP and can rebind to the membrane only after nucleotide exchange with a rate λ. **(b)** MinD binds to flat membrane regions as well and recruits MinD-ATP from the cytosol. Binding to flat regions is, however, less favored due to the lower concentration of MinJ-DivIVA complexes. **(c)** Simulation of the reaction-diffusion model in a 3D rod-shaped cell; shown is the membrane-bound MinD density distribution. As initial condition we take the steady state distribution of the scenario where DivIVA is localized at the poles (left figure). At simulation start we assume that MinD is losing its affinity to the poles by making the recruitment and detachment rate uniform on the entire cell membrane (this is for example the case at the onset of septum formation). From left to right the time evolution of membrane-bound MinD is shown, where the very right side shows the final steady state density distribution. We find that polar localization of MinD becomes unstable and that the proteins preferentially localize at the cell center. (**d**) To test whether MinD can be localized at midplane through MinJ-DivIVA complexes after septum formation, we took the same initial condition as in **(c)** and enhanced recruitment and decreased detachment near midcell, respectively. We find that MinD can sharply localize at the septum.

### Single molecule resolution of the Min system reveals cluster formation

Next, we wanted to test these theoretical predictions concerning a dynamic steady state of MinD proteins experimentally, using single molecule resolution microscopy. In contrast to a stationary bipolar gradient of Min proteins from the cell poles, as described before [3, 8] based on a simple thermodynamic binding of Min proteins to a DivIVA/MinJ template, we expect a dynamic relocalization of Min proteins from the cell pole to the septum. This dynamic steady state would reveal Min components along the entire membrane, including the lateral sites at any time. To achieve the highest possible resolution we used photo-activated light microscopy (PALM). Accordingly, strains expressing Dendra2-MinD (BHF011), MinJ-mNeonGreen (JB40) and DivIVA-PAmCherry (JB37) were utilized. While Dendra2 and PAmCherry are photoswitchable/photactivatable FPs that can be converted from green to red/activated with UV light and are hence well suited for PALM [66], mNeonGreen can be used for PALM because of its innate capability to photo-switch [68]. However, it presents some challenges compared to classical photoactivatable FPs as it cannot be pre-bleached and therefore requires more post-processing for reaching a satisfying artifact-free molecule localizations [73]. Nevertheless, all three strains could be successfully imaged in fixed cells with average precisions of 25 to 30 nm (**Figure 4**).

**Figure 4:**
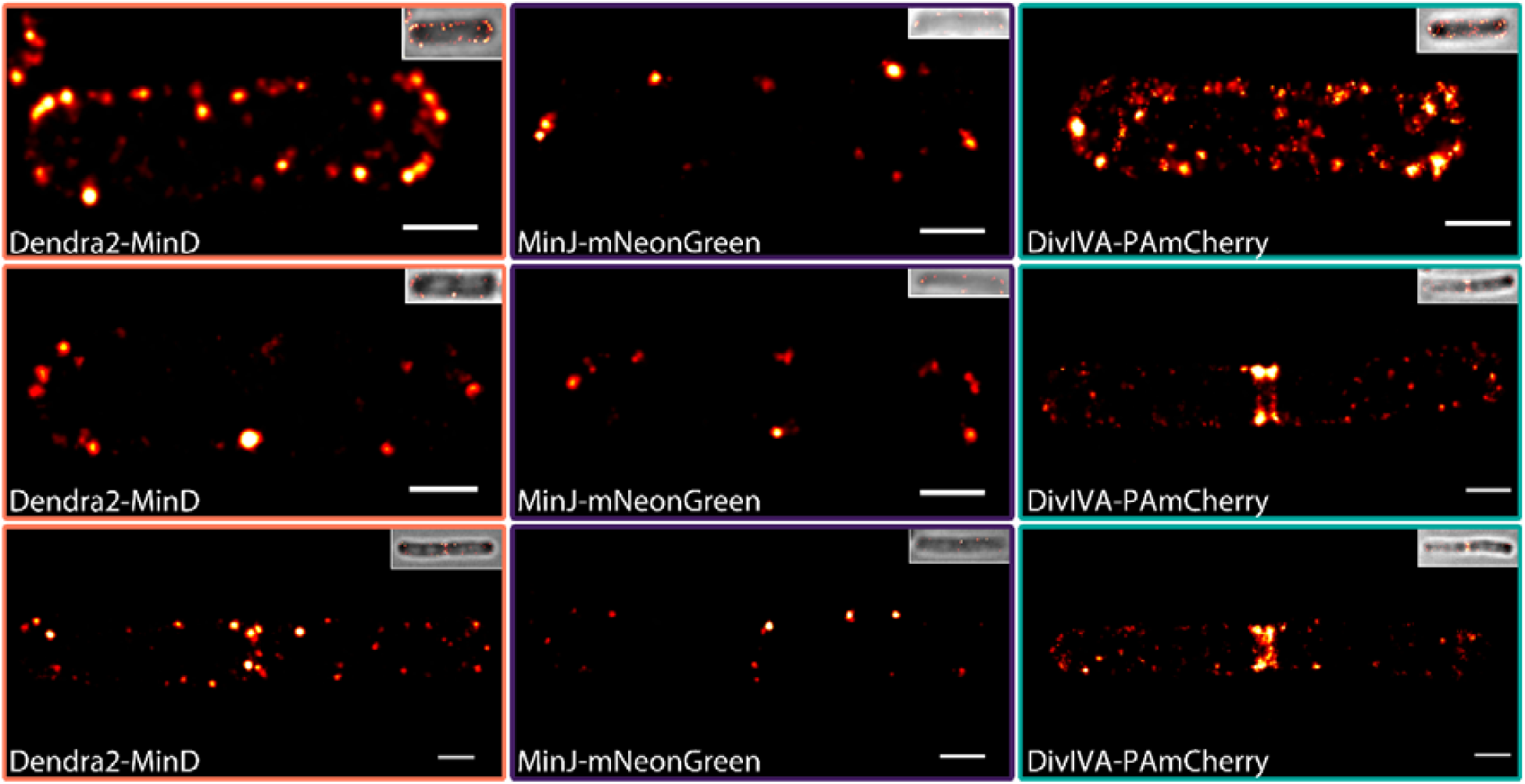
PALM imaging of strains expressing Dendra2-MinD, MinJ-mNeonGreen and DivIVA-PAmCherry. Representative PALM images of Dendra2-MinD (BHF011), MinJ-mNeonGreen (JB40) and DivIVA-PAmCherry (JB37) expressing cells at different divisional states. Upon formation of a division site, DivIVA, MinJ and MinD partially re-localise from the poles to the division septum, where they reside after successful cytokinesis. Samples were fixed prior imaging; every picture represents a different cell. Scale bar 500 nm.

Importantly, we observed all Min proteins not only localized to the cell poles, but also as clusters along the membrane and some apparent cytoplasmic localizations. These protein accumulations can be mainly seen along the membrane for MinJ (**Figure 4 b, middle panel**), while a fraction of MinD and DivIVA could be observed in the cytosol (**Figure 4 b, left & right panel**). The high abundance of these protein accumulations indicates that recruitment of MinD and DivIVA by existing clusters progresses at higher rates compared to individual membrane binding, which is also reflected in the proposed mathematical model. Double rings of MinJ and DivIVA have been reported previously in 3D structured illumination microscopy [75], which could be observed in late divisional cells in PALM as well (**Figure 4 b, middle and bottom panel**). The active enrichment at the young cell pole is consistent with the theoretical model described above and with a role of the Min system in regulation of cell division rather than protection of cell poles from aberrant cell division [61].

To get a deeper insight into the structure and distribution of the imaged proteins and to confirm clustering, a single-molecule point-based cluster analysis was performed for MinD and DivIVA (**Figure 5, Fig. S8, Fig. S9**). Unfortunately, MinJ-mNeonGreen imaging did not produce a sufficient amount of events to be analyzed robustly (**Fig. S8**), as MinJ expression levels are low in comparison and only a small fraction of mNeonGreen molecules blink reliably [73].

**Figure 5:**
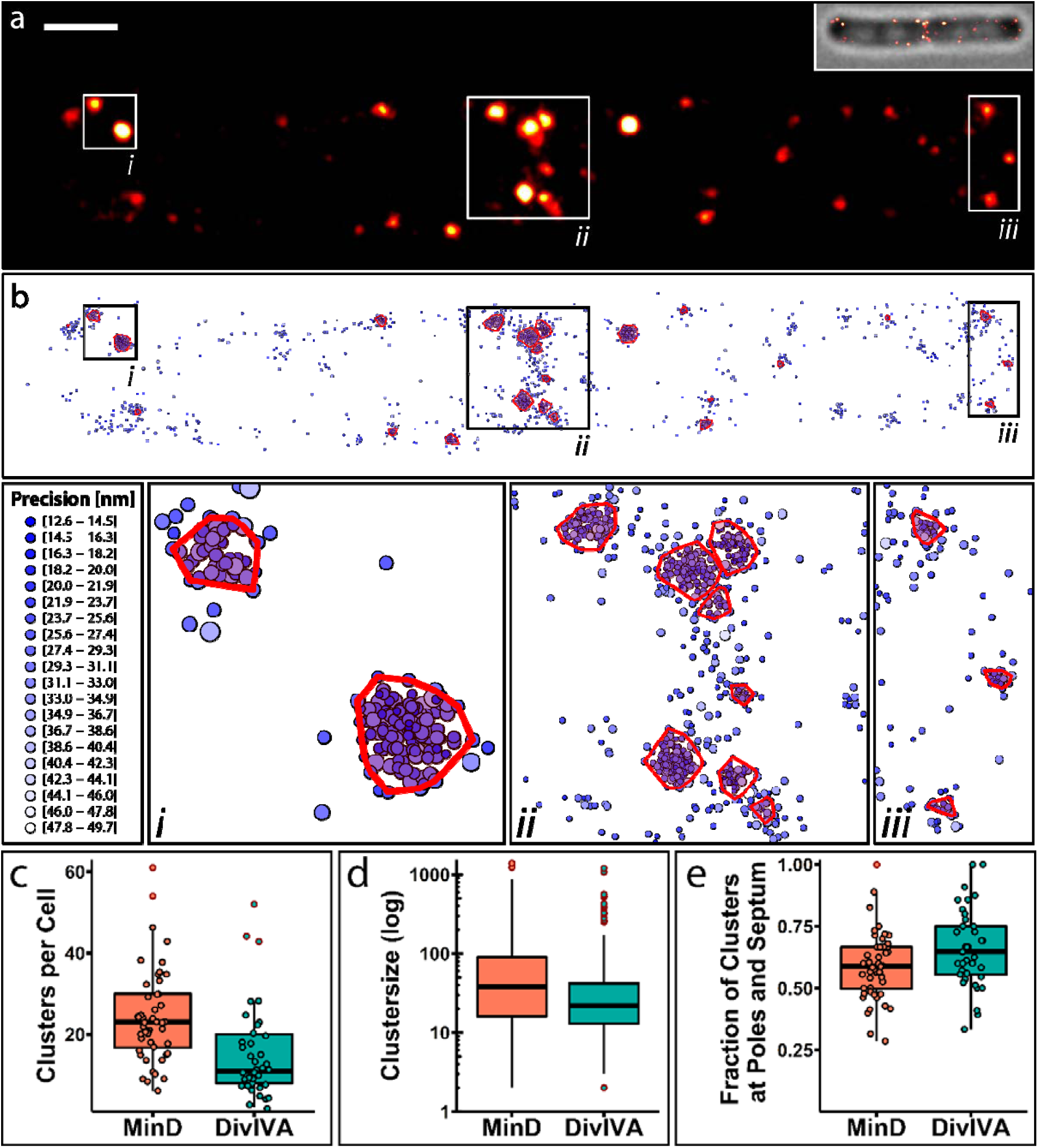
PALM imaging and representative cluster analysis of Dendra2-MinD and DivIVA-PAmCherry. **(a)** Representative PALM image of Dendra2-MinD (BHF011) in a cell in late division state. Scale bar 500 nm. (**b**) Cluster analysis of the same PALM data with three highlighted regions **(*i, ii*** and ***iii*)**. Cluster analysis was performed in R using the OPTICS algorithm from the DBSCAN package. Every point indicates a single event and thus a Dendra2-MinD/DivIVA-PAmCherry protein, precision is indicated by color and size of the circle **(c)** Boxplot of the number of clusters of Dendra2-MinD and DivIVA-PAmCherry per cell (MinD, n_cells_ = 48, DivIVA, n_cells_ =37). **(d)** Boxplot of the number of proteins per cluster, no jitter is shown due to high sample number (Dendra2-MinD, n_clusters_ = 1171, DivIVA-PAmCherry, n_clusters_ = 586). **(e)** Boxplot of fraction of clusters localized at poles and septa per cell (MinD, n_cells_ = 48, DivIVA, n_cells_ =37). Outliers in boxplots are indicated by red outline.

In total, we recorded 151,887 events in 48 cells for Dendra2-MinD, while 52,377 events of DivIVA-PAmCherry could be recorded in 37 cells, respectively. When identifying clusters with at least 10 molecules per cluster, 55.61% (84,470) of all Dendra2-MinD events were organized in clusters, while 52.27% (27,379) events of DivIVA-PAmCherry could be assigned to clusters. Thus, the average prevalence of clusters per cell was higher for MinD (24 clusters per cell) when compared to DivIVA (15 clusters per cell) (**Figure 5 c**). The size of these clusters varied greatly (**Figure 5 d**), an average size of 72 MinD proteins per cluster was determined, while the average number of DivIVA proteins per cluster was 47. However, we also observed some very large clusters with up to 1390 MinD proteins and 1198 DivIVA proteins, respectively. Analysis of the relative position of all clusters per cell revealed a high tendency for clusters to form around poles and septa (**Figure 5 e**), where around two-thirds of DivIVA clusters (66%) and more than half of MinD clusters (59%) were observed, while the rest was found along the lateral membrane or the cytosol. This correlates well with the idea that most of MinD is recruited to negative membrane curvature (poles and septum) by DivIVA via MinJ. MinD also binds to flat membrane areas where it recruits more MinD from the cytosol. This is less favored due to the lower concentration of MinJ-DivIVA complexes, which is reflected in our simulations and data. Our data also reveal that MinD and DivIVA seem to accumulate, and cytosolic proteins therefore have a higher tendency to bind to existing clusters than to free membrane surfaces. We did not observe a large proportion of MinD dimers and also no homogeneous binding of MinD or DivIVA to the membrane.

## Discussion

The Min reaction network has been extensively studied in various organisms [8, 81]. In *E. coli*, it was found to be a highly dynamic and self-organizing system capable of pole-to-pole oscillation, a prime example for intracellular protein pattern formation [34]. The two core components in this network, MinE and MinD, cycle between membrane and cytosol and are sufficient to induce robust protein patterns both *in vivo* and *in vitro* [17, 18, 27, 82, 83]. Therefore, it has been puzzling that the Min system in *B. subtilis* was described to form a rather stationary bipolar gradient from poles to midcell, although MinC and MinD are well conserved. The differences are mainly accredited to the absence of MinE that stimulates ATP hydrolysis and thus membrane detachment of MinD. Instead, the curvature-sensing DivIVA was found to recruit MinCDJ to the negatively curved poles. However, MinC has been shown to dynamically relocalize to midcell prior division in fluorescence microscopy [65], and the same study highlights open questions in the current Min model for *B. subtilis*, pointing out that earlier studies were conducted using strains that artificially overexpress Min network components, thereby possibly masking dynamic populations.

In this study, we analyzed protein dynamics of the *B. subtilis* Min system based on experiments conducted with native expression levels of fluorescently labeled Min components. Firstly, we found all components to be highly dynamic. MinD had the shortest recovery time of the three investigated proteins, while MinJ and DivIVA were both considerably slower compared to MinD, but in a similar range when compared to each other (Tab. 2). Similar tendencies were detected when comparing mobile and immobile fractions, where MinD had the highest mobile fraction with almost 80 % of the protein taking part in the recovery. With diffusion coefficients between 0.057 µm^2^/s and 0.0034 µm^2^/s, the three proteins were found in an expected range for membrane (-associating) proteins in bacteria [84]. Considering the nature of DivIVA, which binds to the membrane and stabilizes itself at negative curvature, and MinJ as an integral membrane protein, it is not surprising that the cytosolic MinD is around 10 fold faster in recovery. This observation leads to the speculation of a relatively fast exchange of membrane bound MinD proteins at the division septum, considering relative high total protein quantities (**Tab. S1**) in combination with a large mobile fraction and fast recovery when bleaching these sites. DivIVA total protein numbers were found to be around half of MinD, while MinJ was by far the least abundant Min component. These findings correlate with the corresponding fluorescence intensity and appearance when imaging the respective Min proteins tagged with the same fluorophore during mid-exponential growth (e.g. **Fig. S2**).

Moreover, knocking out single or multiple components had impact on the dynamics of the respective upstream recruiting factor, validating interaction between MinD and MinJ, and MinJ and DivIVA, respectively that were observed in genetic studies before [58, 59]. Based on this interaction network and the respective protein behaviors in combination with the knowledge gained from the *E. coli* Min system, a mathematical model was designed.

We propose a minimal reaction-diffusion model that correctly reproduces qualitative features of MinD localization in *B. subtilis*. We extracted the parameters for the model from our measurements (protein numbers and diffusion coefficients, **Tab. S1** and **Tab. 2**) and from previous work on intracellular protein pattern formation [30, 32, 34, 79]. The basic assumption of the model is that DivIVA acts as a spatial template for MinJ and MinD, which we accounted for implicitly through a space-dependent recruitment and detachment rate for MinD. From the computational analysis of the model (FEM simulations) we found that localization of MinD to the poles or the division site corresponds to a dynamic equilibrium state of the reaction-diffusion equation. Further, our results show that a geometric effect alone is sufficient to guide MinD to the division site, therefore highlighting the importance of realistic 3D models.

Our model can be straightforwardly extended to include the explicit dynamics of DivIVA and MinJ. As the exact reaction network of the Min system in *B. subtilis* remains elusive, a theoretical model could help in identifying the essential components of Min dynamics. Following the same approach as for the *E. coli* Min system, reconstitution of the Min system *in vitro* would also help to dissect the complexity of the system and to make the comparison between experiments and theory even more feasible. We believe that our theoretical approach may serve as a basis for future studies addressing protein dynamics in *B. subtilis*.

We note that the observed dynamics are not compatible with a division-site selection system, because ongoing division is needed for correct localization and dynamics of the Min system in *B. subtilis*. This is in line with data obtained by Elizabeth Harry’s lab that deletion of Min proteins does not abolish midcell positioning of the Z-ring in *B. subtilis* [85] and our own data describing reduced disassembly of Z-rings in absence of the Min system [61]. The model we propose includes a yet unknown protein or mechanism that stimulates MinD-ATP hydrolysis. The uniform hydrolysis rate *k*_*H*_ in our model was predicted to be similar to the closely related MinD in *E. coli*, which is stimulated by MinE [23, 27]. The responsible protein or mechanism has yet to be elucidated, but the presence of cytosolic and membrane-bound MinD fractions and their respective dynamics as well as the well-conserved ATPase domain argue very convincingly for its existence.

Additionally, we investigated the Min components with single-molecule resolution, revealing a strong tendency for cluster formation and these clusters are also found at the lateral sides of the cell membrane. The lateral Min assemblies have not been resolved by conventional light microscopy images and hence the idea of an exclusive polar Min assembly manifested. Clusters of MinD and DivIVA are indeed frequently observed close to poles and midcell. In accordance with the mathematical model, we hence hypothesize that a fraction of MinD will diffuse away from these primary binding sites after recruitment. Most of this fraction will quickly unbind the membrane due to the lack of stabilization, and will be recruited again by DivIVA-MinJ to either pole or septum clusters. Proteins that are part of a cluster will show less exchange or dynamic behavior, further decreasing towards the center, as it can be typically observed in protein clusters [86]. This mechanism could tightly regulate spatio-temporal localization of MinCD and, likewise, aid in transitioning from polar localization to septal localization rapidly upon septum formation, as DivIVA and MinJ would transition to the septum first. Since the current view on the task of the Min system in *B. subtilis* proposes a role downstream of cell division, all components need to be concentrated at the septum in time to inhibit a second round of division by promoting the disassembly of the division apparatus [61].

## Acknowledgments

This research was supported by the Deutsche Forschungsgemeinschaft within the Transregio Collaborative Research Center (TRR 174) “Spatiotemporal Dynamics of Bacterial Cells” (project P03 to E.F. and P05 to M.B.).We thank the members of the M.B. lab for discussions, feedback and comments on the manuscript.

## Author Contributions

M.B. and E.F. conceived the study, H.F. constructed the strains, performed the in vivo experiments incl. microscopy and analyzed the data, L.W. and E.F. developed the theoretical model and performed the mathematical analysis. All authors wrote the manuscript.

## Material and Methods

### Bacterial strains, plasmids and oligonucleotides

The oligonucleotides, plasmids and strains used in this study are listed in **Tab. 3, Tab. 4** and **Tab. 5**, respectively. *E. coli NEB* Turbo was used to amplify and maintain plasmids.

**Tab. 3:**
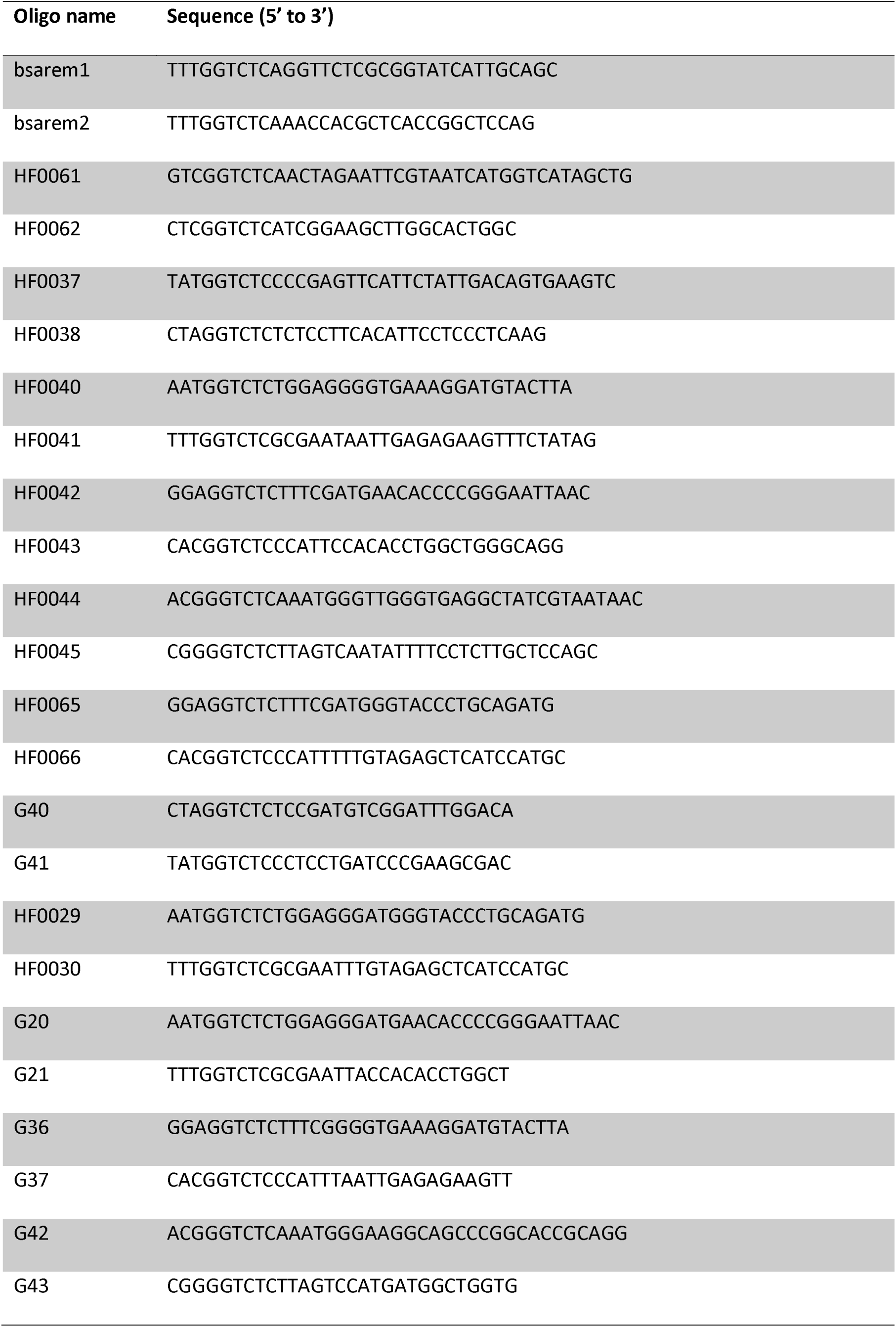
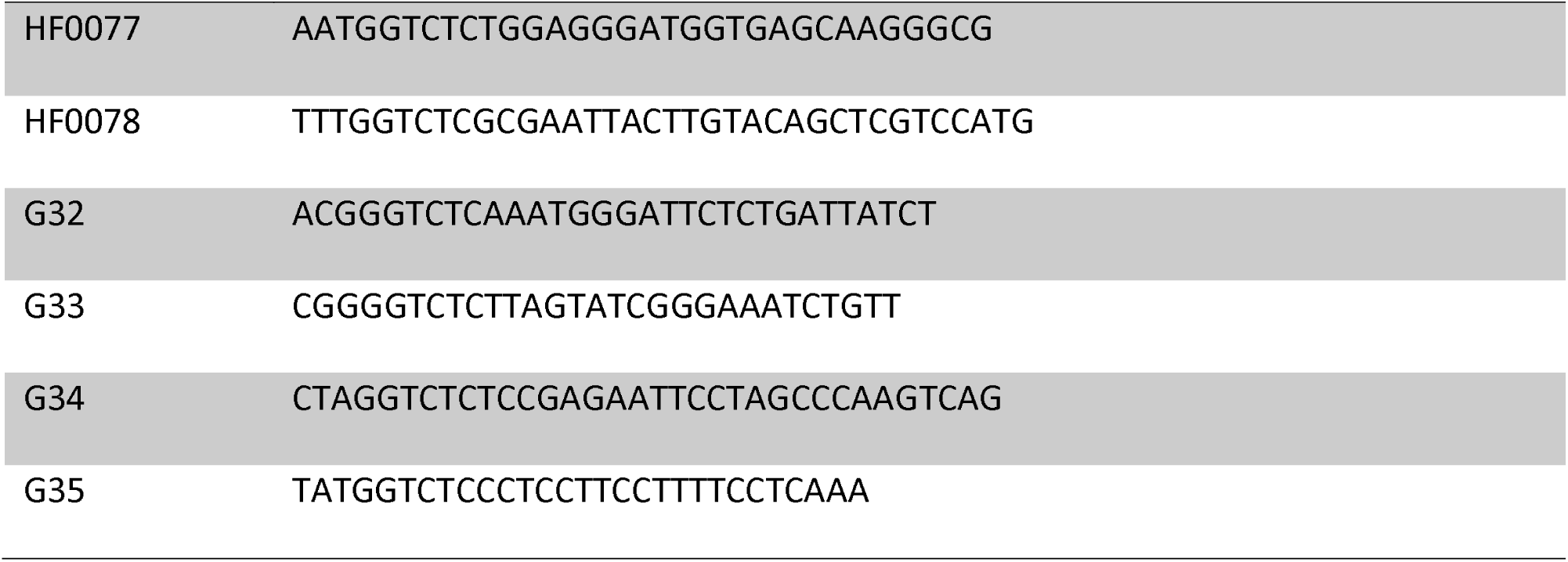
Oligonucleotides used in this study.

### Strain construction

Golden Gate assembly

Fragments for Golden Gate assembly were amplified from *B. subtilis* 168 (trpC2) genomic DNA or template plasmids via PCR with the respective primers containing directional overhangs (**Tab. 3**). The vector pUC18mut was also amplified via PCR to introduce BsaI restriction sites and allow subsequent digestion of circular PCR template with DpnI, which only cuts methylated DNA. Plasmid construction was verified via individual control digestion and DNA sequencing. Correct plasmids were transformed into *B. subtilis* 168 with the respective genetic background (**Tab. 5**) and selected for the introduced resistance (**Tab. 4**). Resistant candidates were confirmed with PCR and microscopy.

**Tab. 4:**
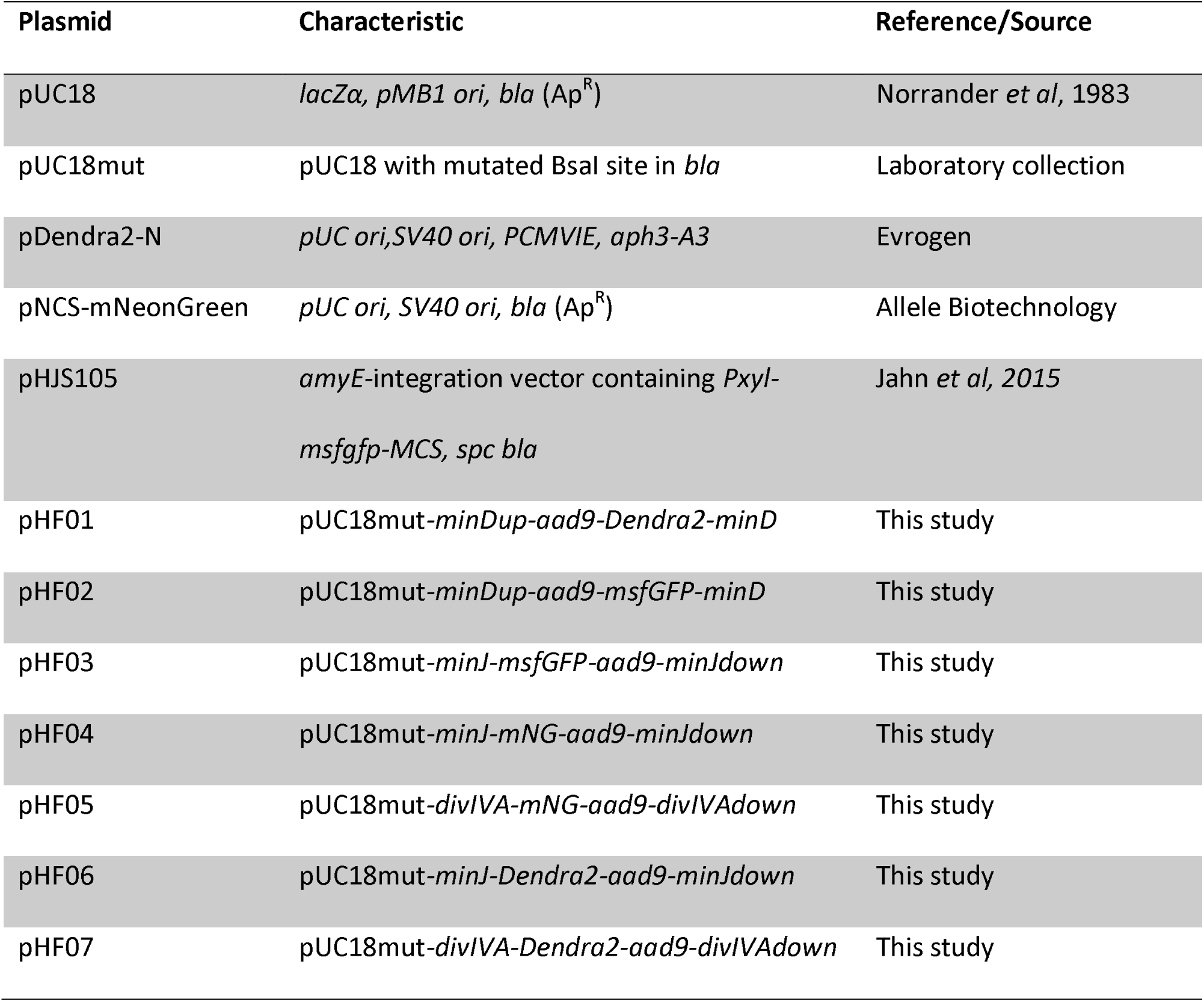
Plasmids used in this study.

**Tab. 5:**
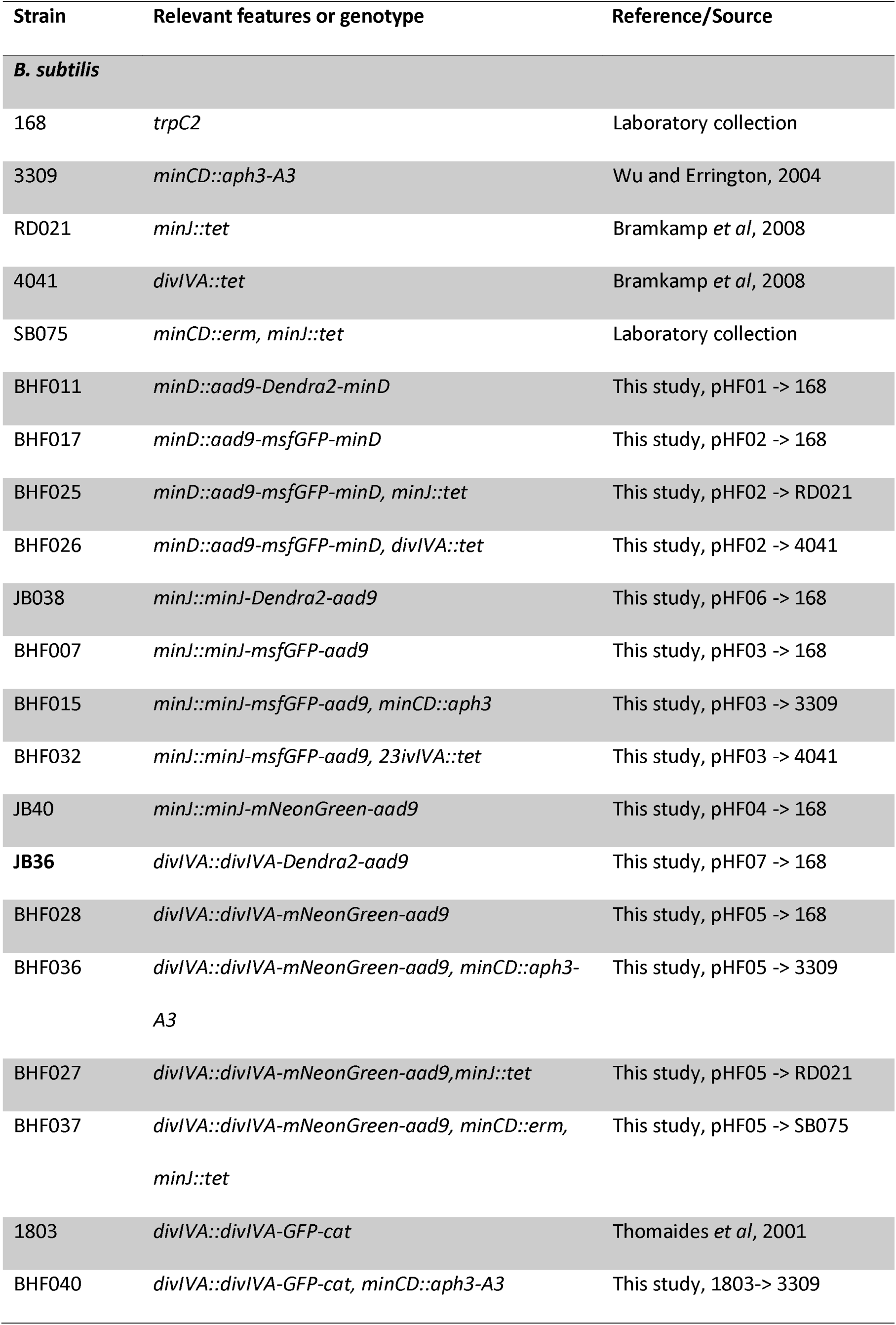
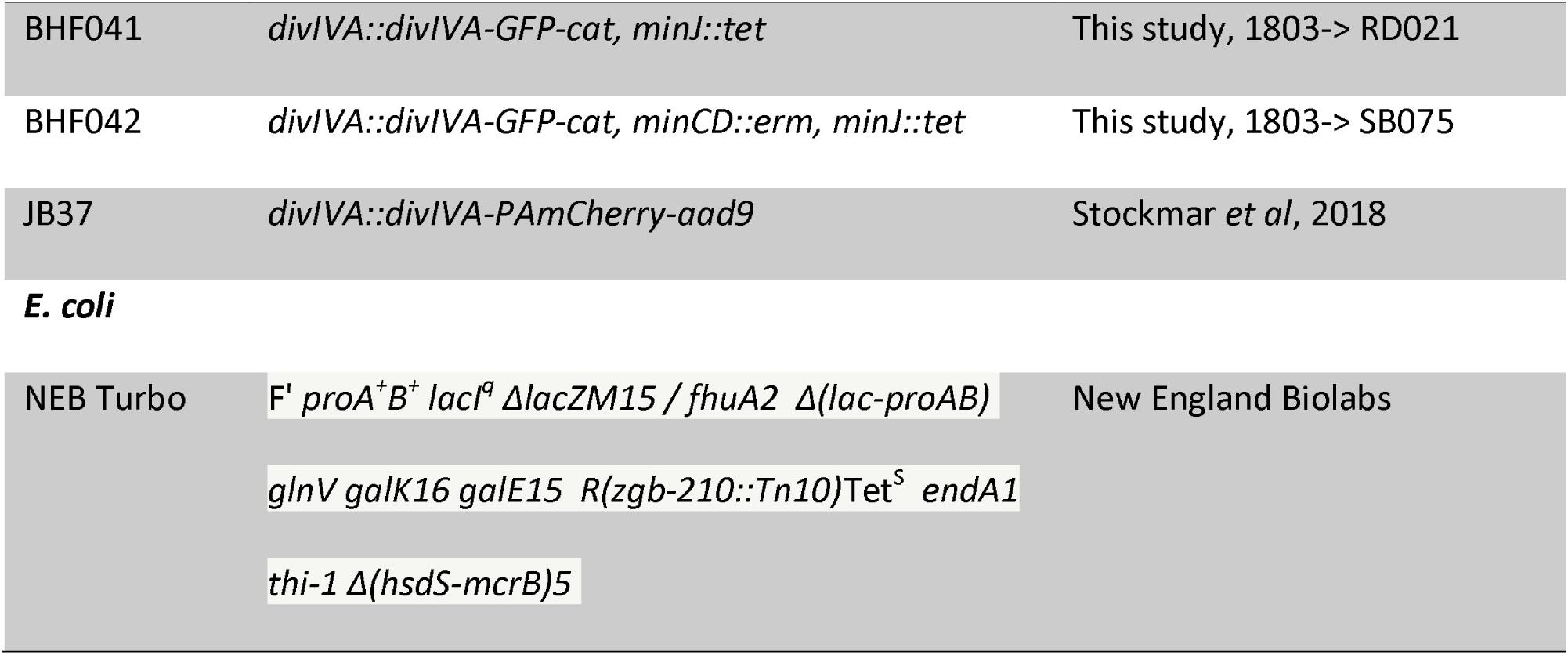
Strains used in this study.

| Strain | Relevant features or genotype | Reference/Source |
| --- | --- | --- |
| ***B. subtilis*** |  |  |
| 168 | *trpC2* | Laboratory collection |
| 3309 | *minCD::aph3-A3* | Wu and Errington, 2004 |
| RD021 | *minJ::tet* | Bramkamp *et al*, 2008 |
| 4041 | *divIVA::tet* | Bramkamp *et al*, 2008 |
| SB075 | *minCD::erm, minJ::tet* | Laboratory collection |
| BHF011 | *minD::aad9-Dendra2-minD* | This study, pHF01 -> 168 |
| BHF017 | *minD::aad9-msfGFP-minD* | This study, pHF02 -> 168 |
| BHF025 | *minD::aad9-msfGFP-minD, minJ::tet* | This study, pHF02 -> RD021 |
| BHF026 | *minD::aad9-msfGFP-minD, divIVA::tet* | This study, pHF02 -> 4041 |
| JB038 | *minJ::minJ-Dendra2-aad9* | This study, pHF06 -> 168 |
| BHF007 | *minJ::minJ-msfGFP-aad9* | This study, pHF03 -> 168 |
| BHF015 | *minJ::minJ-msfGFP-aad9, minCD::aph3* | This study, pHF03 -> 3309 |
| BHF032 | *minJ::minJ-msfGFP-aad9, 23ivIVA::tet* | This study, pHF03 -> 4041 |
| JB40 | *minJ::minJ-mNeonGreen-aad9* | This study, pHF04 -> 168 |
| **JB36** | *divIVA::divIVA-Dendra2-aad9* | This study, pHF07 -> 168 |
| BHF028 | *divIVA::divIVA-mNeonGreen-aad9* | This study, pHF05 -> 168 |
| BHF036 | *divIVA::divIVA-mNeonGreen-aad9, minCD::aph3-A3* | This study, pHF05 -> 3309 |
| BHF027 | *divIVA::divIVA-mNeonGreen-aad9, minJ::tet* | This study, pHF05 -> RD021 |
| BHF037 | *divIVA::divIVA-mNeonGreen-aad9, minCD::erm, minJ::tet* | This study, pHF05 -> SB075 |
| 1803 | *divIVA::divIVA-GFP-cat* | Thomaides *et al*, 2001 |
| BHF040 | *divIVA::divIVA-GFP-cat, minCD::aph3-A3* | This study, 1803-> 3309 |
| BHF041 | *divIVA::divIVA-GFP-cat, minJ::tet* | This study, 1803-> RD021 |
| BHF042 | *divIVA::divIVA-GFP-cat, minCD::erm, minJ::tet* | This study, 1803-> SB075 |
| JB37 | *divIVA::divIVA-PAmCherry-aad9* | Stockmar *et al*, 2018 |
| ***E. coli*** |  |  |
| NEB Turbo | F' *proA<sup>+</sup>B<sup>+</sup> lacI<sup>q</sup> ΔlacZM15 / fhuA2 Δ(lac-proAB)*<br>*glnV galK16 galE15 R(zgb-210::Tn10)Tet<sup>S</sup> endA1*<br>*thi-1 Δ(hsdS-mcrB)5* | New England Biolabs |

pHF01 [pUC18mut-*minDup-aad9-Dendra2-minD*] was created by a Golden Gate assembly of 5 fragments: 1) PCR with primers HF0061 and HF0062 with pUC18mut as template (yielding a linear pUC18mut; 2) PCR with primers HF0037 and HF0038 and 168 genomic DNA (containing the region upstream of *minD*); 3) PCR with primers HF0040 and HF0041 and JB40 genomic DNA (containing the spectinomycin adenyltransferase *aad9*); 4) PCR with primers HF0042 and HF0043 and pDendra2-N plasmid DNA (containing the *Dendra2 gene*); 5) PCR with primers HF0044 and HF0045 and 168 genomic DNA (containing the N-terminal region of *minD*).

pHF02 [*pUC18mut-minDup-aad9-msfGFP-minD*] was created by a Golden Gate assembly of 5 fragments: 1) PCR with primers HF0061 and HF0062 with pUC18mut as template (yielding a linear pUC18mut; 2) PCR with primers HF0037 and HF0038 and 168 genomic DNA (containing the region upstream of *minD*); 3) PCR with primers HF0040 and HF0041 and JB40 genomic DNA (containing the spectinomycin adenyltransferase *aad9*); 4) PCR with primers HF0065 and HF0066 and pHJS105 (containing the *msfGFP gene*); 5) PCR with primers HF0044 and HF0045 and 168 genomic DNA (containing the N-terminal region of *minD*).

pHF03 [pUC18mut*-minJ-msfGFP-aad9-minJdown*] was created by a Golden Gate assembly of 5 fragments: 1) PCR with primers HF0061 and HF0062 with pUC18mut as template (yielding a linear pUC18mut; 2) PCR with primers G40 and G41 and 168 genomic DNA (containing the C-terminal region of *minJ*); 3) PCR with primers HF0029 and HF0030 and pHJS105 (containing the *msfGFP gene*); 4) PCR with primers G36 and G37 and JB40 genomic DNA (containing the spectinomycin adenyltransferase aad9); 5) PCR with primers G42 and G43 and 168 genomic DNA (containing the region downstream of *minJ*).

pHF04 [pUC18mut-*minJ-mNG-aad9-minJdown*] was created by a Golden Gate assembly of 5 fragments: 1) PCR with primers HF0061 and HF0062 with pUC18mut as template (yielding a linear pUC18mut; 2) PCR with primers G40 and G41 and 168 genomic DNA (containing the C-terminal region of *minJ*); 3) PCR with primers HF0077 and HF0078 and pNCS-mNeonGreen DNA (containing the *mNeonGreen gene*); 4) PCR with primers G36 and G37 and JB40 genomic DNA (containing the spectinomycin adenyltransferase aad9); 5) PCR with primers G42 and G43 and 168 genomic DNA (containing the region downstream of *minJ*).

pHF05 [pUC18mut-*divIVA-mNG-aad9-divIVAdown*] was created by a Golden Gate assembly of 5 fragments: 1) PCR with primers HF0061 and HF0062 with pUC18mut as template (yielding a linear pUC18mut; 2) PCR with primers G34 and G35 and 168 genomic DNA (containing the C-terminal region of *divIVA*); 3) PCR with primers HF0077 and HF0078 and pNCS-mNeonGreen DNA (containing the *mNeonGreen gene*); 4) PCR with primers G36 and G37 and JB40 genomic DNA (containing the spectinomycin adenyltransferase aad9); 5) PCR with primers G32 and G33 and 168 genomic DNA (containing the region downstream of *divIVA*).

pHF06 [pUC18mut-*minJ-Dendra2-aad9-minJdown*] was created by a Golden Gate assembly of 5 fragments: 1) PCR with primers HF0061 and HF0062 with pUC18mut as template (yielding a linear pUC18mut; 2) PCR with primers G40 and G41 and 168 genomic DNA (containing the C-terminal region of *minJ*); 3) PCR with primers G20 and G21 and pDendra2-N plasmid DNA (containing the *Dendra2 gene*); 4) PCR with primers G36 and G37 and JB40 genomic DNA (containing the spectinomycin adenyltransferase *aad9*); 5) PCR with primers G42 and G43 and 168 genomic DNA (containing the region downstream of *minJ*).

pHF07 [pUC18mut-*divIVA-Dendra2-aad9-divIVAdown*] was created by a Golden Gate assembly of 5 fragments: 1) PCR with primers HF0061 and HF0062 with pUC18mut as template (yielding a linear pUC18mut; 2) PCR with primers G34 and G35 and 168 genomic DNA (containing the C-terminal region of *divIVA*); 3) PCR with primers G20 and G21 and pDendra2-N plasmid DNA (containing the *Dendra2 gene*); 4) PCR with primers G36 and G37 and JB40 genomic DNA (containing the spectinomycin adenyltransferase *aad9*); 5) PCR with primers G32 and G33 and 168 genomic DNA (containing the region downstream of *divIVA*).

### Media and growth conditions

*B. subtilis* was grown on nutrient agar plates using commercial nutrient broth and 1.5% (w/v) Agar at 37°C overnight. To reduce inhibitory effects, antibiotics were only used for transformations and when indicated, since allelic replacement is stable after integration (chloramphenicol 5 µg ml^−1^, tetracycline 10 µg ml^−1^, kanamycin 5 µg ml^−1^, spectinomycin 100 µg ml^−1^, erythromycin 1 µg ml^−1^).

For growth curves, *B. subtilis* was inoculated to an OD_600_ 0.05 from a fresh overnight culture and grown in LB (lysogeny broth) [10 g l^−1^ tryptone, 10 g l^−1^ NaCl and 5 g l^−1^ yeast extract] at 37°C with aeration in baffled shaking flasks (200 rpm) to OD_600_ 1. Subsequently, cultures were diluted to OD_600_ 0.1 in fresh LB and measured every hour for at least 6 hours.

For microscopy, *B. subtilis* was inoculated to an OD_600_ 0.05 from a fresh overnight culture and grown in MD medium – a modified version of Spizizen Minimal Medium [87] – at 37°C with aeration in baffled shaking flasks (200 rpm) to OD_600_ 1. MD medium contains 10.7 mg ml^−1^ K_2_ HPO_4_, 6 mg ml^−1^ K_2_H_4_ PO, 1 mg ml^−1^ Na_3_ citrate, 20 mg ml^−1^ glucose, 20 mg ml^−1^ L-tryptophan, 20 mg ml^−1^ ferric ammonium citrate, 25 mg ml^−1^ L-aspartate and 0.36 mg ml^−1^ MgSO_4_ and was always supplemented with 1 mg ml^−1^ casamino acids. Subsequently, cultures were diluted to OD_600_ 0.1 in fresh MD medium and grown to OD_600_ 0.5 (exponential phase).

For epifluorescence and time-lapse imaging (e.g. FRAP), *B. subtilis* cells were mounted on pre-warmed 1.5% MD agarose pads, sealed with paraffin and incubated 10 min at 37°C before microscopic analysis. When used, FM4-64 dye was added to the agarose pad before polymerization (1 µM final).

For PALM imaging, a 0.5 ml portion of *B. subtilis* cells were fixed by addition of formaldehyde (1.5% (w/v) final concentration) and incubated for 20 min at 37°C. Subsequently, cells were washed (1 min, 5000 rpm), resuspended in fresh MD supplemented with 10 mM glycine to stop the crosslinking reaction and incubated for 10 min at 37°C. Then, cells were washed 2 more times with MD containing 10mM glycine. In a final washing step, the pellet was resuspended in 50 µl of MD with 10 mM glycine to reach higher cell density. Cells were mounted on chambered coverslips (µ-slide 8 well, Ibidi) containing 200 µl MD with 10 mM glycine, which were pretreated for 30-60 min with 0.1% Poly-L-lysine and successively washed 3 times with MD containing 10 mM glycine. Furthermore, TetraSpeck microspheres (100 nm, ThermoFisher) were added in a dilution that results in about 3-10 beads per field of view. To help sedimentation of cells and beads and to reach a uniform attachment to the glass surface, the chambered coverslip was centrifuged at 4000 rpm for 10 min in a bucket-swing rotor (Eppendorf).

### Typhoon imaging and western blots

To confirm the presence of full-length protein fusions and for quantitative analysis, *B. subtilis* strains were inoculated from an overnight culture to OD_600_ 0.05 in the morning, and grown to OD_600_ 0.5 in 10 ml LB medium (MD for quantitative studies) at 37°C. Cells were then diluted 1/10 and grown again to mid-exponential phase (OD_600_ 0.5). Cultures were centrifuged at 13.000 rpm for 1 minute, washed once with lysis buffer (10 mM Tris, pH 7.5, 150 mM NaCl, 500 µM EDTA, 1 mM PMSF), and resuspended in lysis buffer with additional 10 mg/ml lysozyme (Sigma-Aldrich), 10 μg/ml DNAse I (Roche) and 100 μg/ml RNAse A (Roche), concentrating the sample to OD_600_ 30. After incubation at 37°C for 20 minutes, the sample was briefly vortexed to crack remaining intact cells. 30 µl of sample was then mixed with 10 µl of 4x SDS-PAGE loading buffer (200 mM Tris-HCL pH6.8, 400 mM DTT. 8% SDS. 0,4% bromophenol blue and 40% glycerol). For typhoon imaging and subsequent western blotting, samples were incubated for either 20 min at room temperature, while some samples meant exclusively for western blotting were incubated at 95°C for 10 minutes for full denaturation (indicated in **Fig. S4**) 10 or 20 µl of sample were then separated by SDS Page in 12% Bis-Tris gels. For visualization of green fluorescent fusions, gels were imaged in a Typhoon Trio (GE Healthcare; PMT 600-800, Excitation 488 nm, Emission 526 SP). For western blotting, proteins were blotted onto 0.2 µm polyvinylidene difluoride (PVDF) membranes. Proteins were visualized via anti-mCherry (polyclonal), anti-mNG (monoclonal) or anti-Dendra (polyclonal) antibodies, respectively.

To quantify Dendra2-fusions of MinD, MinJ and DivIVA via in-gel fluorescence, three biological triplicates were prepared and imaged as described above, while avoiding oversaturation. The total number of MinD molecules was taken from a publication that utilized targeted mass spectrometry to determine absolute protein amounts of *B. subtilis* at mid-exponential phase in minimal medium with glucose [76]. Relative quantification was then performed using ImageJ by measuring and comparing intensity of the bands.

### Fluorescence Microscopy

For strain characterization, microscopy images were taken on a Zeiss Axio Observer Z1 microscope equipped with a Hamamatsu OrcaR^2^ camera using a Plan-Apochromat 100×/1.4 Oil Ph3 objective (Zeiss). Dendra2, GFP, msfGFP and mNeonGreen fluorescence was visualized with filterset 38 HE eGFP shift free (Zeiss) and FM4-64 membrane dye was visualized with filterset 63 HE mCherry (Zeiss). The microscope was equipped with an environmental chamber set to 37°C. Digital images were acquired with Zen software (Zeiss).

For FRAP experiments a Delta Vision Elite (GE Healthcare, Applied Precision) equipped with an Insight SSI™ illumination, an X4 laser module and a CoolSnap HQ2 CCD camera was used. Images were taken with a 100×oil PSF U-Plan S-Apo 1.4 NA objective. A four color standard set InsightSSI unit with following excitation wavelengths (DAPI 390/18 nm FITC 475/28 nm, TRITC 542/27 nm, Cy5 632/22 nm); single band pass emission wavelengths (DAPI 435/48 nm, FITC 525/48 nm, TRITC 597/45 nm, Cy5 679/34 nm) and a suitable polychroic for DAPI/FITC/TRITC/Cy5 were used. GFP, msfGFP and mNeonGreen were visualized using FITC settings and exposure times between 0.1 s (msfGFP, GFP) and 0.2 s (mNeonGreen). Bleaching was performed using a 488 nm laser (50 mW) with 10% power and a 0.005 – 0.01 s pulse. Frequency of acquisition and total amount of images were chosen according to the individual recovery times after initial testing with various settings.

Analysis of the images was performed using ImageJ 1.51 s. The corrected total cell fluorescence (CTCF) was calculated according to following formula: CTCF = Integrated Density—(Area of selected cell X Mean fluorescence of unspecific background readings) [88]. For FRAP experiments unspecific background was subtracted for every ROI (see above). The CTCF of the septa was divided by the CTCF of the whole cell to account for photobleaching during acquisition. The respective quotient of the unbleached spot was always set as 1 for normalization. Since *B. subtilis* keeps growing during time-lapse experiments like FRAP, the bleached spot moves in the field of view as cells elongate. Therefore, a macro in Fiji was created to dynamically follow and center the bleached spot through the frames of acquisition without any bias, which resulted in more precise FRAP curves. To determine half-time recovery and mobile/immobile fractions, the FRAP curve from the normalized recovery values was fitted to an exponential equation:

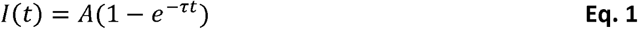

where *I*(*t*) is the normalized FRAP curve, A the final value of the recovery, *τ* the fitted parameter and *t* the time after the bleaching event. After determination of the fitted coefficients, they can be used to determine mobile (*A*) and immobile (1-*A*) fractions, while following equation was used to determine halftime recovery (**Eq. 2**):

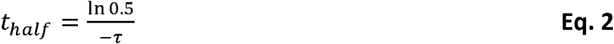

where *t*_*half*_ is the halftime recovery and *τ*the fitted parameter. Diffusion coefficients were then calculated with the following formula:

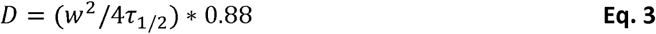

according to Axelrod et al., 1976 [89] – where *D* is the diffusion coefficient, *w* is the radius of the circular laser beam and *τ*_1/2_ is the time when fluorescence recovery reaches half height of total recovery. To estimate the bleaching spot radius, cells expressing cytosolic GFP were fixed with 1.5% formaldehyde (v/v) as described above, mounted on agarose pads, bleached at laser powers of 10% to 100% in increments of 10% and imaged right after bleaching. The radius was measured in ImageJ and averaged per triplicate to calculate the function of bleach radius over laser power. Graphs and statistics were created in R 3.3.1 [90] utilizing the packages ggplot2 [91] and nlstools [92].

### Reaction-diffusion equations

The setup of our mathematical model is based on previous approaches for intracellular protein dynamics [30, 32, 34, 79]. Specifically, we present a minimal model to account for DivIVA mediated MinD localization. The model includes the following set of biochemical reactions: *(i)* attachment of MinD-ATP (with volume concentration *u*_*DT*_) from the bulk to the membrane with constant rate *k*_*D*_; *(ii)* recruitment of bulk MinD-ATP to the membrane by membrane-bound MinD (with areal concentration *u*_*d*_) with rate *k*_*dD*_; *(iii)* hydrolysis and detachment of membrane-bound MinD into bulk MinD-ADP (*u*_*DD*_) with rate *k*_*H*_; *(iv)* reactivation of bulk MinD-ADP by nucleotide exchange to MinD-ATP with rate *λ*. The system of ensuing reaction–diffusion equations reads

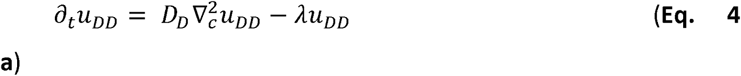

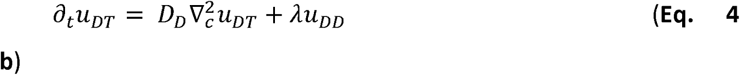

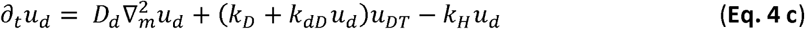

where the subscript *c* or *m* denotes that the nabla operator acts in the bulk or on the membrane, respectively. These equations are coupled through nonlinear reactive boundary conditions at the membrane surface stating that the biochemical reactions involving both membrane-bound and bulk proteins equal the diffusive flux onto and off the membrane:

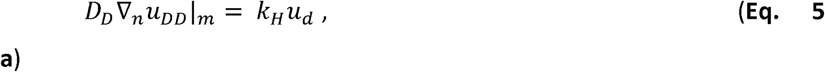

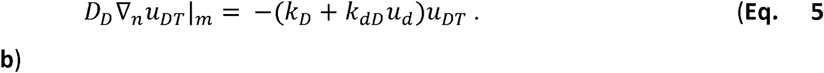

Here, the subscript *n* denotes that we take the nabla operator acting along the outward normal vector of the boundary (membrane). The set of reaction–diffusion equations conserve the total mass of MinD. Hence, the total particle number *N*_*D*_ of MinD obeys the relation

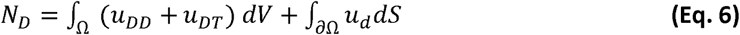

We simulated the set of reaction–diffusion equations in a spherocylindrical geometry in three-dimensional space (3D) using the Finite-Element software *COMSOL v5.3a*; for an illustration of the geometry used see **Fig. S7**. The length and height were set to typical values known for *B. subtilis* cells, *L* = 2.8 *µm* and *h* = *0*.*85 µm*, respectively. The mean total density of MinD was set to *MinD*] = 2450 *µm*^−3^ for all simulations (see **Tab. S2**) We assume that, in addition to MinD self-recruitment, MinJ recruits MinD-ATP from the bulk to the membrane and that membrane-bound MinD is stabilized by DivIVA-MinJ complexes on the membrane. We model the interaction of MinD with MinJ and DivIVA implicitly through space-dependent recruitment and detachment rates. To this end, we assume that the recruitment rate is amplified by a factor *α* and that the detachment rate is reduced by a factor *β* at regions of high negative curvature (such as the poles or the septum). This assumption is motivated by experiments which suggest that MinD localization is dependent on MinJ and that DivIVA acts as a scaffold which stabilizes MinJ and MinD (see main text). We therefore set the recruitment and detachment rate to 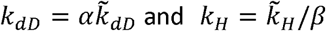 at regions of high negative curvature (**Fig. S7**) where *α* and *β* denotes dimensionless amplification and reduction prefactors, respectively. The parameters 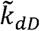 and 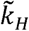 denotes the uniform recruitment and detachment rate which one would obtain if interactions between MinD and MinJ-DivIVA complexes are neglected, i.e. if *α* = *β* = 1 (see below).

### Simulation of the model

#### Polar localization

In a cell with no pre-existing division apparatus, the Min system localizes at the poles of the bacteria (see discussion in main text). We model this case by setting *α* = 4 and *β* = 3 at the polar caps and *α* = *β* = 1 for the remaining part of the rod-shaped geometry (see **Fig. S7 b**) The uniform rates were set to 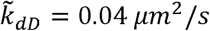 and 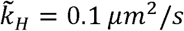 as given in the main text. Simulations show that MinD can be pinned to the cell poles for non-uniform kinetic parameters (**Figure 3 c** left in main text).

#### Depletion of MinD at the poles

Next, we tested if the polar distribution of MinD decays to a homogeneous protein distribution along the membrane when the rates are uniform over the whole cell body. To this end, we used the steady state polar distribution of MinD (as obtained above) as initial condition for a simulation with uniform rates in the entire geometry, i.e. *α* = 1, *β* = 1, respectively. We find that for uniform rates MinD proteins preferentially localize near cell center (**Figure 3 c** left to right in main text). The reason for this unexpected inhomogeneous protein distribution is an interplay between reactions, diffusion and cell geometry. In short, this effect can be explained as follows: MinD detaches from the membrane in an inactive MinD-ADP state and can therefore not rebind to the membrane until it exchanges its nucleotide to switch to an active state MinD-ATP. This results in a source degradation process with the decay length set by 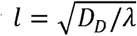. Due to membrane curvature these reaction volumes overlap near the cell poles, which implies an accumulation of inactive MinD-ADP at the cell poles. For a detailed discussion of this geometric effect we refer to [79]. Note that once DivIVA loses its affinity to the cell poles, this effect alone could explain the switch of MinD localization from the poles to midcell. Hence, fully 3D simulations are necessary to reveal the influence of cell geometry on the protein dynamics.

#### Localization at septum

The curvature sensing protein DivIVA targets the division site and guides MinJ and MinD to the septum (see discussion in the main text). Above we showed that MinD localizes to the cell poles if the recruitment and detachment rate of MinD are altered at the poles due to interactions with MinJ and DivIVA. For uniform rates, however, the MinD density distribution is spread around mid cell but not sharply localized at the septum as observed in experiments. Sharp localization of MinD at mid cell requires interaction with DivIVA and MinJ and we therefore model this case in the same way as for polar localization: First, we define a narrow region with width *s*_*w*_ = 0.14 *µm* at mid cell which represents the septum (**Fig. S7 c**). We set again *α* = 4 and *β* = 3 at this geometric region to model the interactions of MinD with MinJ and DivIVA implicitly through a modified recruitment and detachment rate. Simulations of the model show that MinD localizes sharply at the septum (**Figure 3 d** left to right in main text).

### Photo-activated localization microscopy (PALM) and cluster analysis

PALM imaging was performed with the microscope system ELYRA P.1 (Zeiss) and the accompanied Zen software. It is equipped with a 405 nm Diode-Laser (50 mW), a 488 nm laser (200 mW), a 561 nm laser (200 mW) and a 640 nm laser (150 mW). Furthermore, an alpha Plan-Apochromat 100x/1,46 Oil DIC M27 objective (Zeiss) was used, in combination with a 1.6x Optovar. The filter sets were the following: a 77 HE GFP+mRFP+Alexa 633 shift free (EX TBP 483+564+642, BS TFT 506+582+659, EM TBP 526+601+688), a 49 DAPI shift free (EX G 365, BS FT 395, EM BP 445/50), a BP 420-480 / LP 750, a BP 495-550 / LP 750, a LP 570 and a LP 655 filter set. Images were recorded with the Andor EM-CCD camera iXon DU 897. Samples expressing mNeonGreen were illuminated with the 488 nm laser at 7.4 mW. Samples expressing Dendra2 or PAmCherry were illuminated with the excitation laser (561 nm, 5.3 mW) and activation laser (405 nm). To avoid co-occurrence of multiple events in the same spot, the power of the activation laser was increased stepwise from 0.008 mW to 1.6 mW. MinJ-mNeonGreen was illuminated in pseudo-TIRF (total internal reflection fluorescence) mode and recorded at 20 Hz with 200 camera gain, while Dendra2-MinD and DivIVA-PAmCherry were imaged with the same camera settings in regular wide-field. Analysis was performed in the Zen Black (Zeiss) software. Detection of single emitters was performed with a peak mask size of 9 pixels and a minimum peak intensity to noise ratio of 6.0., overlapping emitters were discarded. Localization was extrapolated via a 2D Gaussian fitting, and images were drift corrected utilizing a fiducial-based mode with at least 3 beads in focus. Filtering was used to minimize noise, background, out of focus emitters and to exclude beads from the evaluation, according to **Tab. 6**, different for each respective fluorophore.

**Tab. 6:** Filter parameters for PALM imaging of the different strains. Filters were chosen according to the fluorophore behavior in PALM to eliminate background and signal of fluorescent beads from the results.

| Strain/FP | Filter: PSF width at half maximum | Filter: Photon Number |
| --- | --- | --- |
| Dendra2-MinD | 70 - 160 | 70 – 250 |
| MinJ-mNeonGreen | 70 - 160 | 70 – 300 |
| DivIVA-PAmCherry | 60 - 170 | 50 – 500 |

Cluster analysis was performed in R 3.3.1 [90] utilizing the DBSCAN package [93, 94] including OPTICS [95]. Clusters were determined by applying the OPTICS algorithm to the respective molecule tables generated via PALM. The minimal number of points that define a cluster (*minPts*) was defined as 10, reflecting apparent clusters seen in rendered PALM imaging, and a minimum distance between cluster edge points (*epsCl*) of 20 and 30 nm for MinD and DivIVA, respectively, according to the observed density of protein localization.

