## Supplemental Material for "Dynamics of the *Bacillus subtilis* Min system"

### Supplementary Tables

Table S1: Relative quantification of Min proteins fused to Dendra2. Relative amounts of protein were determined via in-gel fluorescence of biological triplicates from cell lysates (see Figure S5). Absolute protein quantities were determined relative to MinD that was quantified in another publication [75] in similar conditions. Values are shown with standard deviation.

| Protein | Relative amount | Total copies per cell |
|---|---|---|
| **MinD** | 100% ± 2.51% | **3544 ± 89** |
| **MinJ** | 16.25% ± 4.36% | **576 ± 25** |
| **DivIVA** | 47.70% ± 3.51% | **1690 ± 59** |

**Table S2: Kinetic rate constants for the MinD dynamics.** The membrane diffusion coefficient, MinD protein density, cell length and cell width are chosen in accordance with our experimental data. The bulk diffusion coefficient, attachment rate, hydrolysis rate and nucleotide exchange rate were estimated from previous approaches for intracellular protein dynamics.

| **Parameter** | **Symbol** | **Value** |
|---|---|---|
| **Bulk Diffusion** | $D_D$ | $16\ \mu m^2 \cdot s^{-1}$ |
| **Membrane Diffusion** | $D_d$ | $0.06\ \mu m^2 \cdot s^{-1}$ |
| **Mean total density** | $[MinD]$ | $2450\ \mu m^{-3}$ |
| **Attachment rate** | $k_D$ | $0.068\ \mu m \cdot s^{-1}$ |
| **Uniform recruitment rate** | $\tilde{k}_{dD}$ | $0.04\ \mu^2 \cdot s^{-1}$ |
| **Uniform hydrolysis rate** | $\tilde{k}_H$ | $0.1\ s^{-1}$ |
| **Recruitment rate amplification factor** | $\alpha$ | $4$ |
| **Hydrolysis rate reduction factor** | $\beta$ | $3$ |
| **Nucleotide exchange rate** | $\lambda$ | $6\ s^{-1}$ |
| **Cell length** | $L$ | $2.8\ \mu m$ |
| **Cell width** | $h$ | $0.85\ \mu m$ |

### Supplementary Figure legends

**Figure S1: Cartoon of strain construction strategy for allelic replacement in *B. subtilis*.** All strains in this study were created to express fluorophore-fusions from their native promoter to sustain native protein levels. Furthermore, they were tested for functionality**.** **(a)** Construction of plasmids was performed with golden-gate cloning, yielding a plasmid that can directly be transformed into *B. subtilis*. **(b)** After transformation, genes for a fluorophore and an antibiotic resistance cassette are integrated into the genomic locus of interest via homologous recombination

**Figure S2: Microscopic images of a selection of strains used in this study.** Columns from left to right: Phase contrast, membrane dye (FM4-64) shown in red, green fluorescent channel depicting the indicated fluorophore and composite of all three channels. Scale bars 2µm.

**Figure S3: Representative microscopy images of FRAP analysis of DivIVA-mNeonGreen. (a)** DivIVA-mNeonGreen expressed in wild type background (BHF028). Images taken before bleaching the indicated spot with a 488 nm laser pulse, directly after bleaching and after recovery of fluorescence. Scale bars 2 µm. **(b)** Representation of the normalized fluorescence recovery in the green channel over time. $T_{1/2}$ = time when fluorescence recovery reaches half height of total recovery, indicated on the graph with a dashed square. The red line represents measured values, black the fitted values.

**Figure S4: Western blots or in-gel fluorescence of native Min protein fusions.** To control for full-length of fluorescent protein fusions with MinD, MinJ or DivIVA, cell lysates were either fully (96°C for 10 min, **d**) or partially (room temperature for 20 min, **a**, **b**, **c** and **e**) denatured and separated via SDS-PAGE. Protein bands were then visualized either directly via in-gel fluorescence **(c, e)** with excitation and emission at 488/526 nm, respectively, or colorimetrically via western-blot with the indicated antibodies (**a**: polyclonal anti-Dendra2, **b**: monoclonal anti-mNeonGreen, **d**: polyclonal anti-mCherry), respectively.

**Figure S5: Relative quantification of native Dendra2 fusions assayed by in-gel fluorescence of SDS-PAGE gels.** Biological triplicates indicated by top right number **(1-3)**. Lysates of the respective strain were partially denatured with SDS loading dye at room temperature for 20 min, loaded in different relative amounts (left 1x, right 2x) and separated via SDS-PAGE. (**a**) Visualization via Typhoon Trio scanner, with excitation at 488 nm and an emission filter of 526 nm. (**b**) Coomassie stain of the respective image as loading control. Results of quantification can be found in **Table S1**.

**Figure S6: Representative microscopy images of FRAP analysis of Min proteins in different knockout backgrounds.(a)** MinJ-msfGFP expressed in Δ*minCD* background (BHF015), and DivIVA-GFP expressed in Δ*minCD* (BHF040), Δ*minJ* (BHF041) and Δ*minCDJ* (BHF042) backgrounds. Images taken before bleaching the indicated spot with a 488 nm laser pulse, directly after bleaching and after recovery of fluorescence. Scale bars 2 µm. **(b)** Representation of the normalized fluorescence recovery in the green channel over time. $T_{1/2}$ = time when fluorescence recovery reaches half height of total recovery, indicated on the graph with a dashed square. The red line represents measured values, black the fitted values.

**Figure S7: Geometry for the simulation of the model. (a)** Sketch of the simulation geometry (spherocylinder). **(b)** Polar localization is achieved by setting $\alpha = 4$ and $\beta = 3$ at the poles (green area), and $\alpha = \beta = 1$ for the remaining part of the geometry. (**c**) Localization at the septum is achieved by setting $\alpha = 4$ and $\beta = 3$ in a narrow region at mid cell (green) and else $\alpha = \beta = 1$.

**Figure S8: PALM imaging and representative cluster analysis of strain JB40 expressing MinJ-mNeonGreen. (a)** PALM image of MinJ-mNeonGreen in a cell in late division state. Scale bar 500 nm. **(b)** Cluster analysis of the same PALM data with three highlighted regions ***(i, ii*** and ***iii).*** Cluster analysis was performed in R using the OPTICS algorithm from the DBSCAN package. Every point indicates a single event and thus a MinJ-mNeonGreen protein, precision is indicated by color and size of the circle.

**Figure S9: PALM imaging and representative cluster analysis of strain JB37 expressing DivIVA-PAmCherry. (a)** PALM image of DivIVA-PAmCherry in a cell in late division state. Scale bar 500 nm. **(b)** Cluster analysis of the same PALM data with three highlighted regions (***i, ii*** and ***iii***). Cluster analysis was performed in R using the OPTICS algorithm from the DBSCAN package. Every point indicates a single event and thus a DivIVA-PAmCherry protein, precision is indicated by color and size of the circle.

### Supplementary Figures

Figure S1

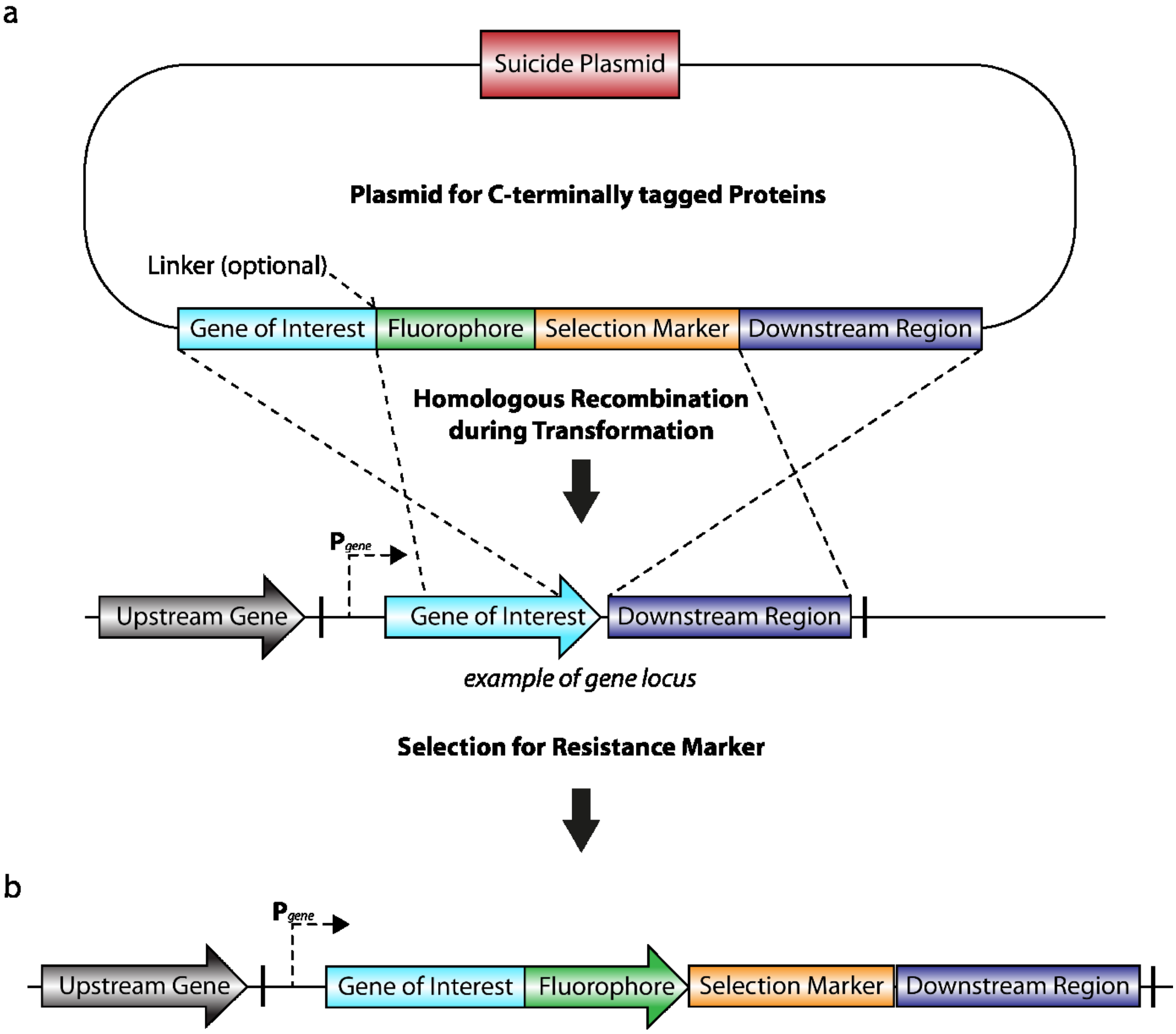

Figure S2

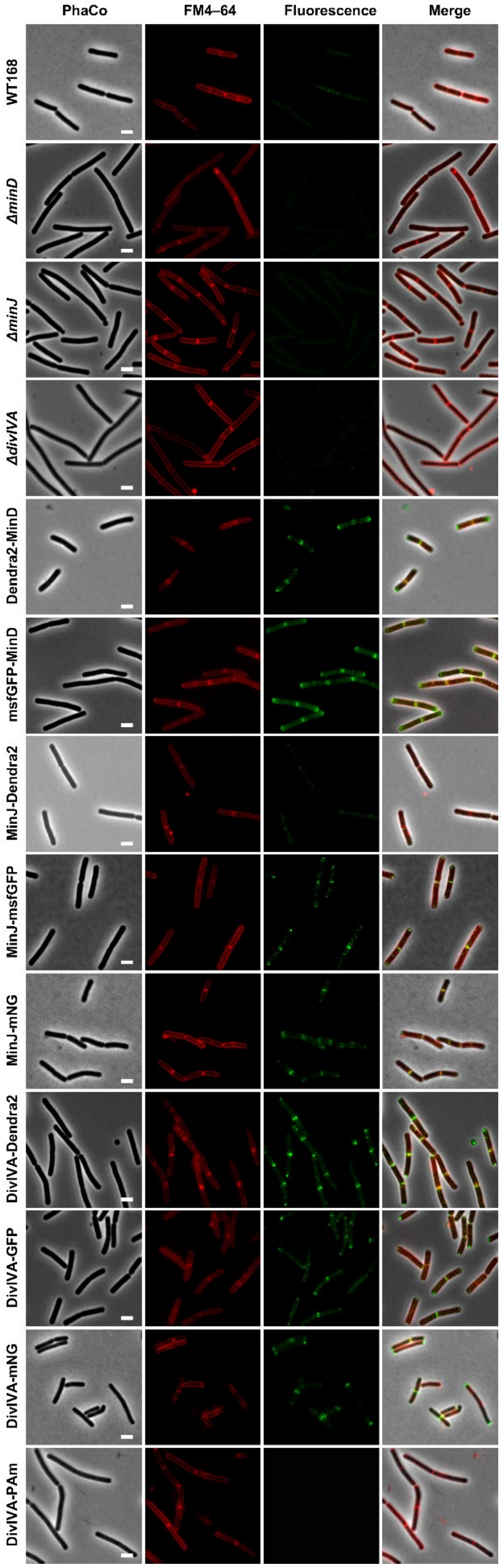
PhaCo
FM4–64
Fluorescence
Merge
WT168
ΔminD
ΔminJ
ΔdivIVA
Dendra2-MinD
msfGFP-MinD
MinJ-Dendra2
MinJ-msfGFP
MinJ-mNG
DivIVA-Dendra2
DivIVA-GFP
DivIVA-mNG
DivIVA-PAm

Figure S3

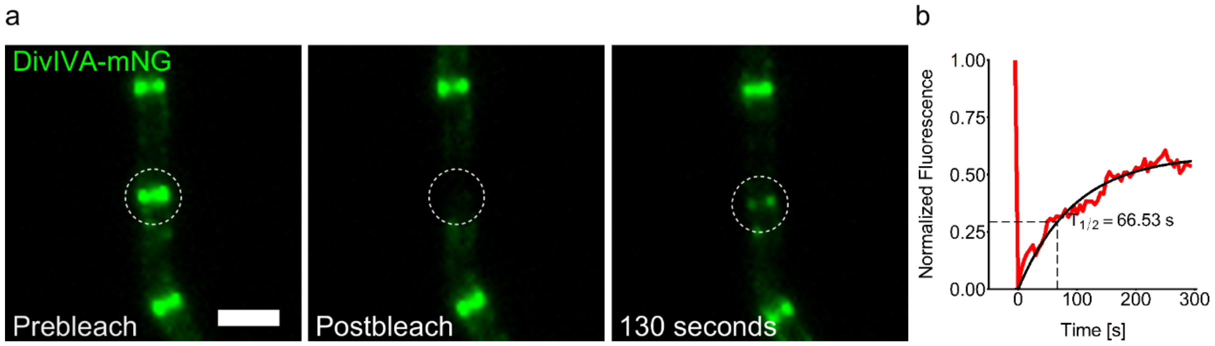
a
DivIVA-mNG
Prebleach
Postbleach
130 seconds
b
1.00
0.75
0.50
0.25
0.00
Normalized Fluorescence
T1/2 = 66.53 s
0
100
200
300
Time [s]

Figure S4

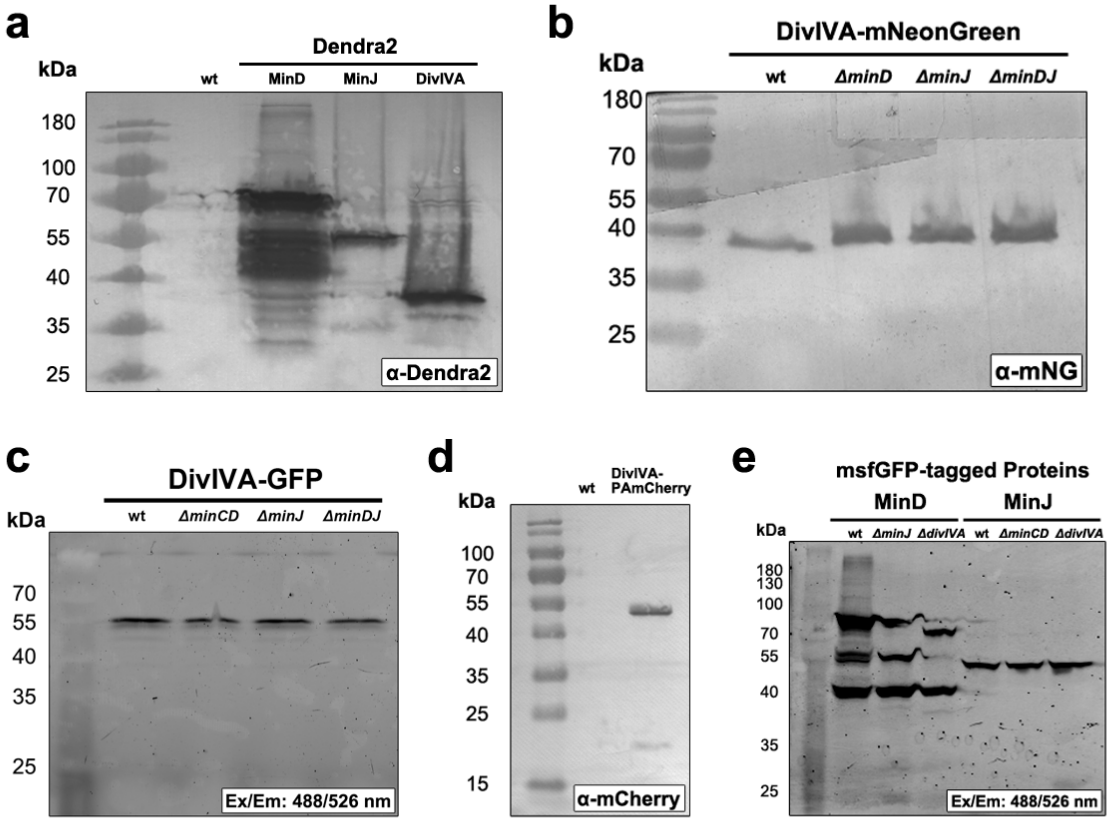
a
kDa
Dendra2
wt
MinD
MinJ
DivIVA
180
100
70
55
40
35
25
α-Dendra2
b
DivIVA-mNeonGreen
kDa
wt
ΔminD
ΔminJ
ΔminDJ
180
70
55
40
35
25
α-mNG
c
DivIVA-GFP
kDa
wt
ΔminCD
ΔminJ
ΔminDJ
70
55
40
35
25
Ex/Em: 488/526 nm
d
kDa
wt
DivIVA-PAmCherry
100
70
55
40
35
25
15
α-mCherry
e
msfGFP-tagged Proteins
MinD
MinJ
kDa
wt
ΔminJ
ΔdivIVA
wt
ΔminCD
ΔdivIVA
180
130
100
70
55
40
35
25
Ex/Em: 488/526 nm

Figure S5

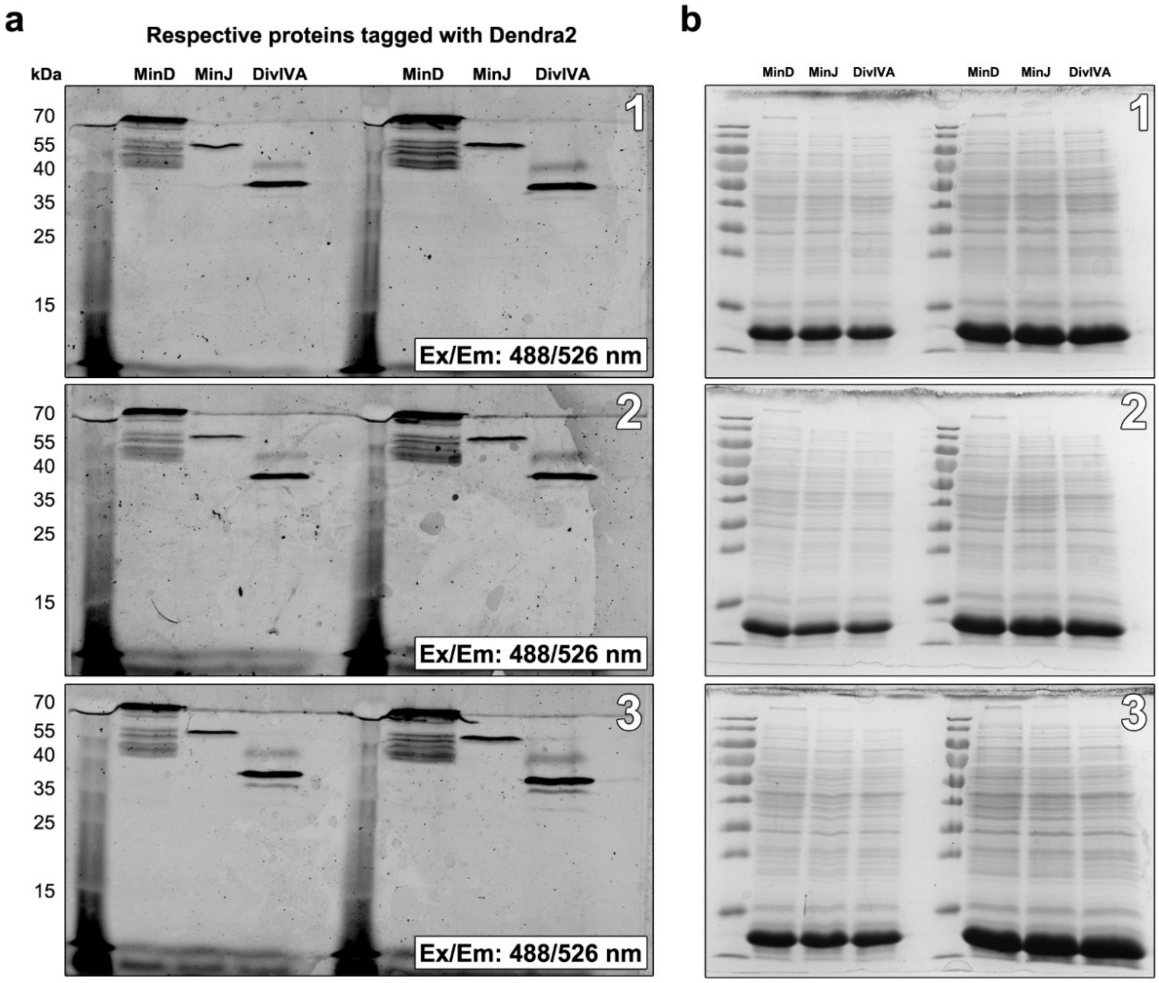
a
Respective proteins tagged with Dendra2
kDa
MinD
MinJ
DivIVA
MinD
MinJ
DivIVA
70
55
40
35
25
15
1
2
3
Ex/Em: 488/526 nm
Ex/Em: 488/526 nm
Ex/Em: 488/526 nm
b
MinD
MinJ
DivIVA
MinD
MinJ
DivIVA
1
2
3

Figure S6

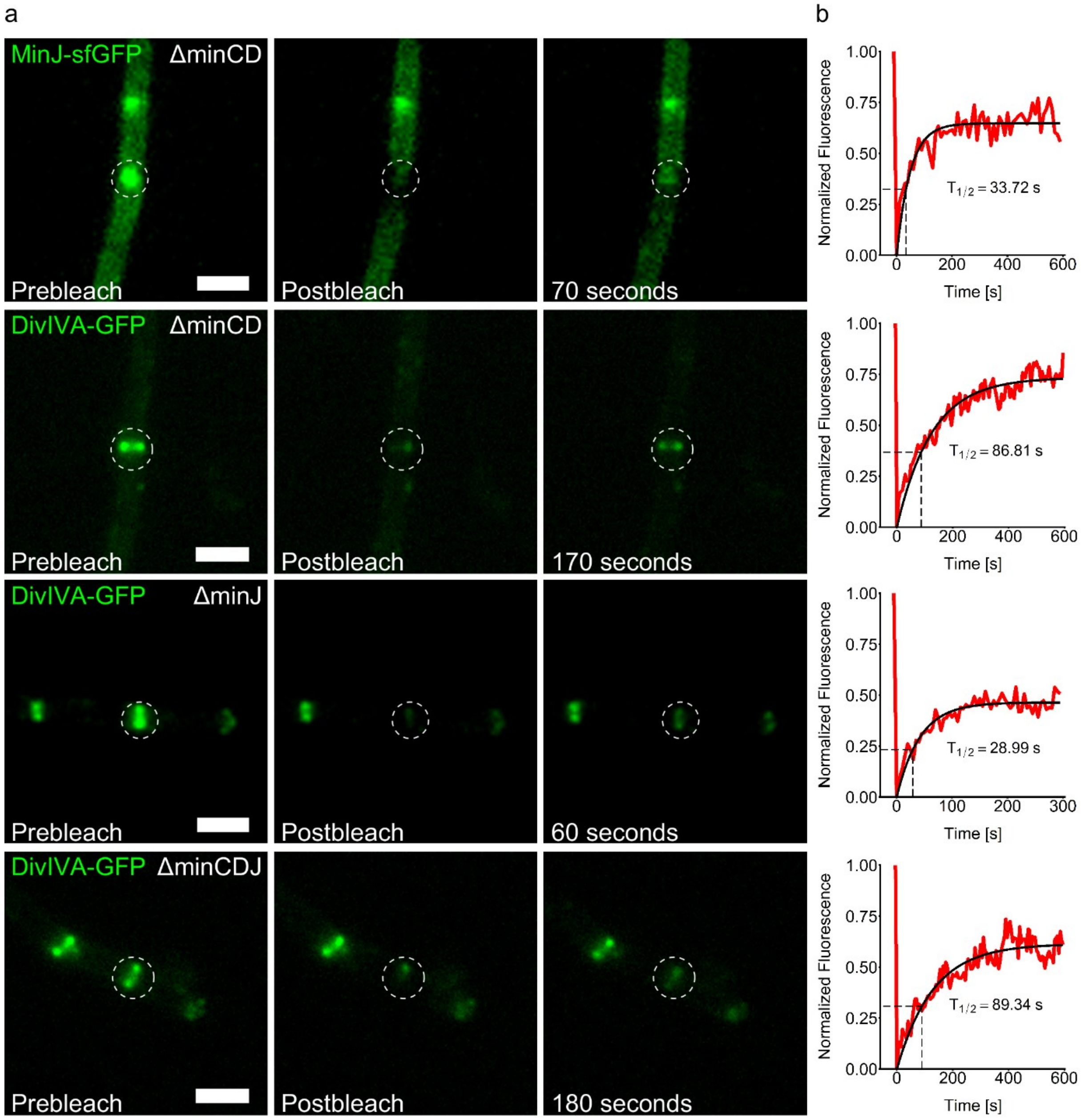

a
MinJ-sfGFP
ΔminCD
Prebleach
Postbleach
70 seconds
DivIVA-GFP
ΔminCD
Prebleach
Postbleach
170 seconds
DivIVA-GFP
ΔminJ
Prebleach
Postbleach
60 seconds
DivIVA-GFP
ΔminCDJ
Prebleach
Postbleach
180 seconds
b
Normalized Fluorescence
Time [s]
T1/2 = 33.72 s
T1/2 = 86.81 s
T1/2 = 28.99 s
T1/2 = 89.34 s

Figure S7

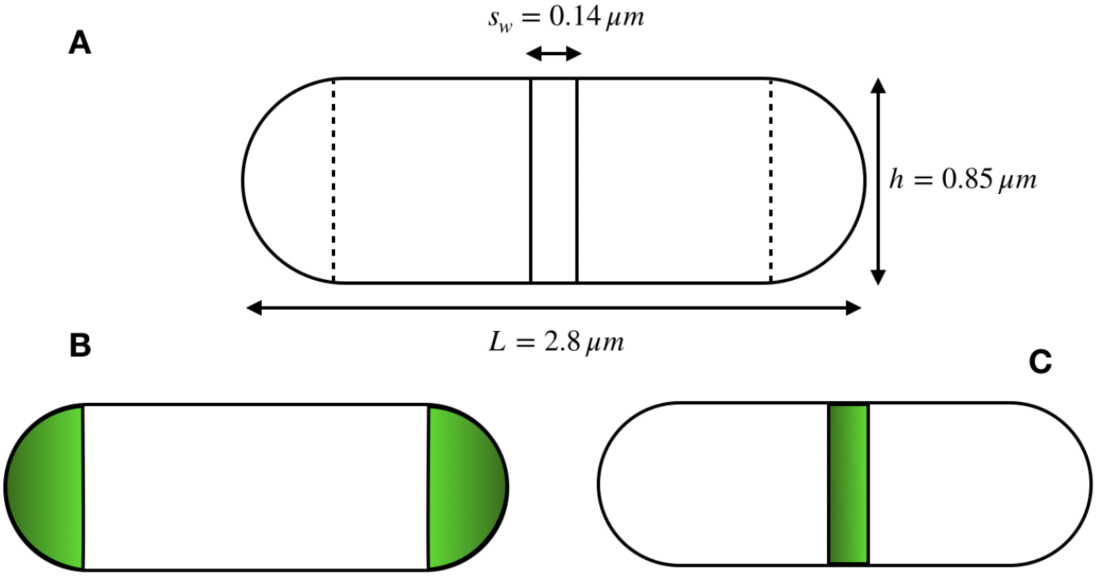

Figure S8

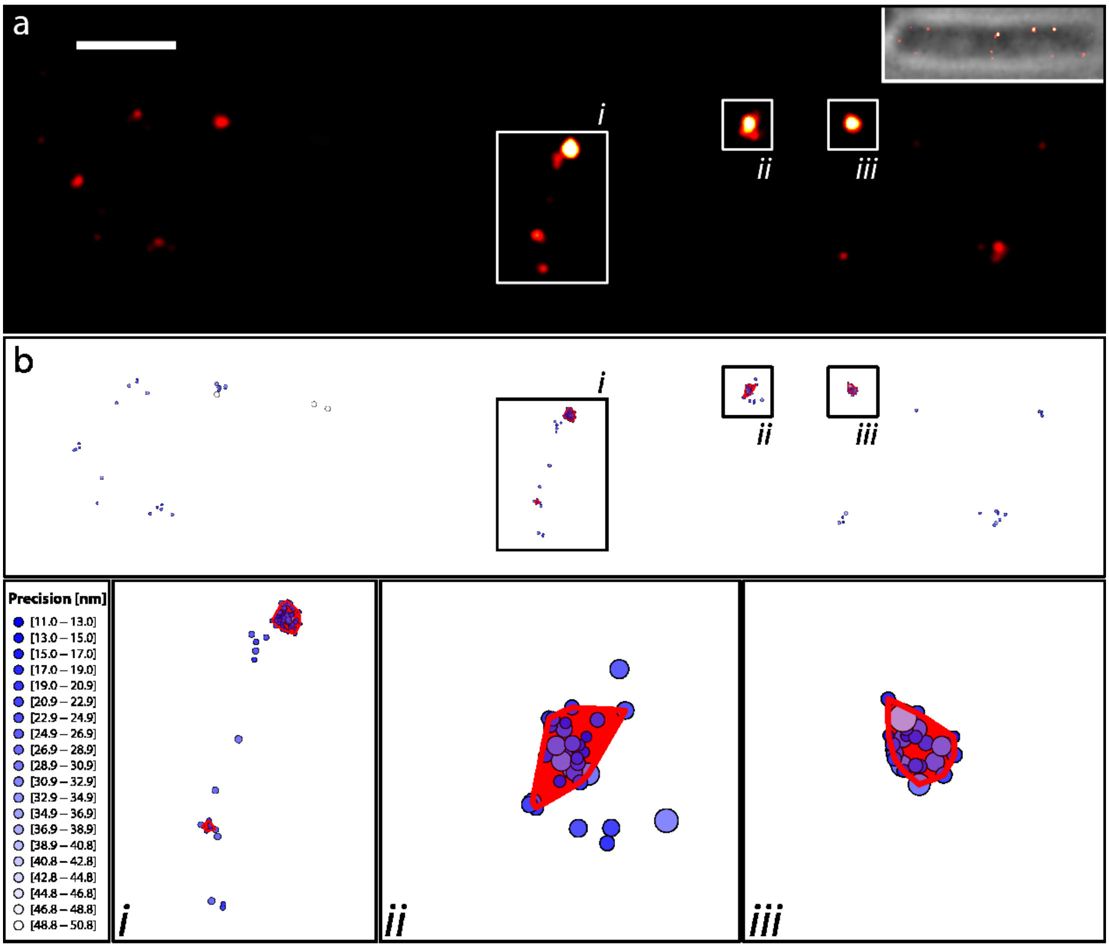
a
i
ii
iii
b
i
ii
iii
Precision [nm]
[11.0 – 13.0]
[13.0 – 15.0]
[15.0 – 17.0]
[17.0 – 19.0]
[19.0 – 20.9]
[20.9 – 22.9]
[22.9 – 24.9]
[24.9 – 26.9]
[26.9 – 28.9]
[28.9 – 30.9]
[30.9 – 32.9]
[32.9 – 34.9]
[34.9 – 36.9]
[36.9 – 38.9]
[38.9 – 40.8]
[40.8 – 42.8]
[42.8 – 44.8]
[44.8 – 46.8]
[46.8 – 48.8]
[48.8 – 50.8]
i
ii
iii

Figure S9

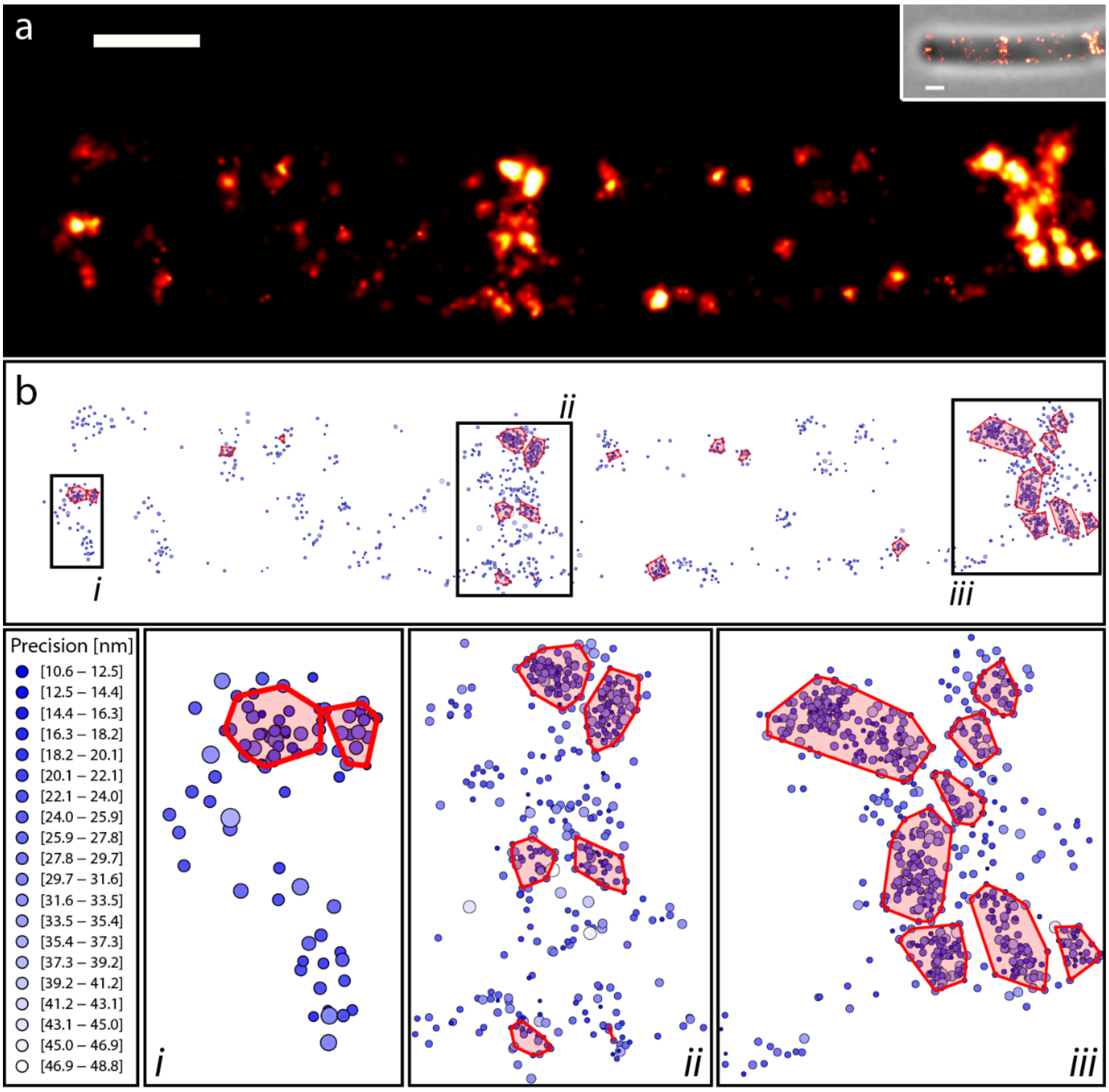

a
b
ii
i
iii
Precision [nm]
[10.6 – 12.5]
[12.5 – 14.4]
[14.4 – 16.3]
[16.3 – 18.2]
[18.2 – 20.1]
[20.1 – 22.1]
[22.1 – 24.0]
[24.0 – 25.9]
[25.9 – 27.8]
[27.8 – 29.7]
[29.7 – 31.6]
[31.6 – 33.5]
[33.5 – 35.4]
[35.4 – 37.3]
[37.3 – 39.2]
[39.2 – 41.2]
[41.2 – 43.1]
[43.1 – 45.0]
[45.0 – 46.9]
[46.9 – 48.8]
i
ii
iii
